# Fear memory-associated synaptic and mitochondrial changes revealed by deep learning-based processing of electron microscopy data

**DOI:** 10.1101/2021.08.05.455246

**Authors:** Jing Liu, Junqian Qi, Xi Chen, Zhenchen Li, Bei Hong, Hongtu Ma, Guoqing Li, Lijun Shen, Danqian Liu, Yu Kong, Qiwei Xie, Hua Han, Yang Yang

## Abstract

Serial section electron microscopy (ssEM) can provide comprehensive 3D ultrastructural information of the brain with exceptional computational cost. Targeted reconstruction of subcellular structures from ssEM datasets is less computationally demanding but still highly informative. We thus developed a Region-CNN-based deep learning method to identify, segment, and reconstruct synapses and mitochondria to explore the structural plasticity of synapses and mitochondria in the auditory cortex of mice subjected to fear conditioning. Upon reconstructing over 135,000 mitochondria and 160,000 synapses, we found that fear conditioning significantly increases the number of mitochondria but decreases their size, and promotes the formation of multi-contact synapses comprising a single axonal bouton and multiple postsynaptic sites from different dendrites. Modeling indicates that such multi-contact configuration increases the information storage capacity of new synapses by over 50%. With high accuracy and speed in reconstruction, our method yields structural and functional insight into cellular plasticity associated with fear learning.

## Introduction

The mammalian brain consists of a vast and complex network of neurons interconnected by specialized sites called synapses. In this network, a neuron may receive input from, and send output to, thousands of other neurons. The concerted activities of neurons, which encode, process, and store information, fundamentally depend on the connectivity patterns of synapses. Thus, it is critical to elucidate the organization of synaptic circuits in order to understand brain functions. Dissecting the synaptic circuit is technically challenging due to small size, complex morphology, dense distribution, and enormous number of synapses. Light microscopy has been used to examine populations of synapses *in vitro* and *in vivo*. Previous work has shown that learning effectively modifies synaptic structures of the mammalian cerebral cortex^1–3^. Auditory fear conditioning (AFC), a common paradigm of associative learning, increases formation of presynaptic boutons and postsynaptic spines in the auditory cortex (A1)^3^. However, although boutons and spines can be visualized using light microscopy, the width of the synaptic cleft is below the diffraction limit and synapses are difficult to discern using light microscopy images^4^. The serial-section electron microscopy (ssEM) technique^5^ overcomes the resolution problem and enables large-scale three-dimensional (3D) reconstruction of brain tissue with nanometer-scale resolution, which is sufficient to resolve the ultrastructural features of synapses, such as presynaptic vesicles, the synaptic cleft, and the postsynaptic density (PSD). However, manual identification and segmentation of synapses from massive ssEM datasets is extremely time-consuming, and thus requires an automated pipeline.

To date, a variety of machine learning-based approaches for synapse detection have been proposed. Some methods require saturated reconstruction or segmentation of neuronal structures prior to synapse detection^6, 7^, which is daunting for large datasets. Some methods do not make full use of the contextual information or structural properties of synapses^8, 9^, making them more prone to errors. Some other methods require nearly isotropic imaging data^10^, a requirement incompatible with standard ssEM, in which the axial resolution (section thickness) is typically much worse than the lateral resolution. Most recently, an indirect method detected synapses by identify the synaptic connectivity (pre- and postsynaptic component of each synapse)^11^. All in all, there is much to be desired in terms of the identification accuracy, speed, and general applicability of automated synapse analysis tools.

Mitochondria play essential roles in cellular functions, such as producing adenosine triphosphate (ATP), and participating in calcium homeostasis^12^. Moreover, synaptic mitochondria are linked to the process of neurotransmitter release and organization of synaptic vesicles^13^. In the past decade, ssEM has been increasingly used to investigate mitochondrial structures. Along with this, a variety of automated methods have been developed to detect mitochondria from ssEM images. One method was based on handcrafted features and traditional classifiers^14–16^, the other on powerful 2D or 3D convolutional neural networks (CNNs)^10, 17^.

In this study, we used Region-CNN (R-CNN) based deep learning algorithms to identify, segment, and reconstruct synapses and mitochondria from ssEM images of the mouse auditory cortex. Our pipeline achieved state-of-the-art accuracy with a speed almost three orders of magnitude faster than human experts, enabling us to sample more than one hundred thousand synapses and mitochondria. Using this method, we studied how a classical learning model, namely AFC, affects the synaptic and mitochondrial organization in the A1 (Figure 1, Extended Data Figure 1). Using automated synapse reconstruction and mathematical modeling, we found that AFC increases multi-synaptic boutons connecting one axonal bouton to multiple different dendrites in the A1, and these 1-to-N connections dramatically increased the information coding capacity, thus representing a synaptic memory engram.

**Figure 1.**
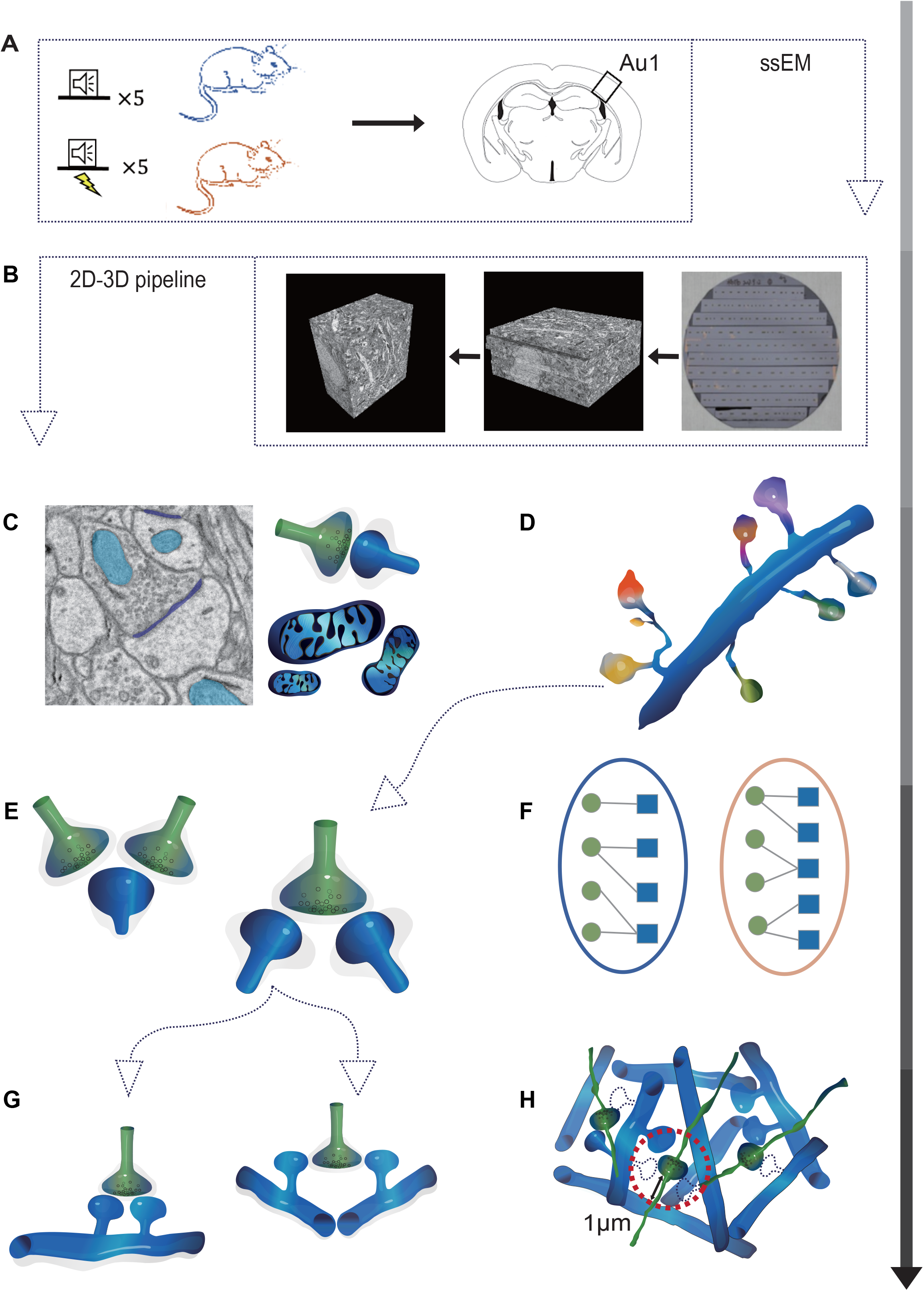
Overall schematic of the experimental procedure. (A) Auditory fear conditioning and sample preparation. Control mice are subjected to 5 repeats of tone pips; fear conditioned mice are subjected to 5 repeats of paired tone pips and foot shocks. At 4 days after conditioning, auditory cortex (A1) tissue blocks are harvested from these mice and prepared for serial section electron microscopy (ssEM). (B) The ssEM image acquisition and alignment procedure. Serial sections are automatically sectioned using automated tape-collecting ultramicrotome (ATUM); these are collected onto 4-inch silicon wafers. The wafers are then imaged using scanning electron microscopy. Raw images are aligned using a SIFT-flow-based, non-linear registration algorithm. (C) Identification of synapses and mitochondria in A1. Synapses and mitochondria are automatically identified using our 2D-3D pipeline based on Region-Convolutional Neural Network (R-CNN). (D) The reconstructed dendrites from the saturated reconstruction by applying the Multicut pipeline^33^. The spine fragments are manually traced to the original dendrite. (E) Multiple-contact synapses (MCSs) localization and classification. MCSs are manually verified and classified into Multi-Synaptic Boutons (MSBs, consisting of a single bouton contacting multiple PSDs) and Multi-synaptic Spines (MSSs, consisting of a single spine contacting multiple boutons) by incorporating the vesicle cloud features. (F) A combinatorial mathematical model is built to simulate the synaptic turnover associated with the fear conditioning process. (G) MSB subtypes. MSBs are classified into single- or multi-dendritic MSBs by incorporating saturated reconstructions to identify the origination dendrites of multiple postsynaptic spines. (H) A synaptic network is built based on ssEM data, using the number of dendrites within a 1-µm radius around a bouton.

## Results

### Auditory fear conditioning as a model for learning and memory

To investigate changes in cellular structures induced by learning and memory in the adult brain, we used a simple and robust behavioral model for associative learning: auditory fear conditioning (AFC). Specifically, conditioned mice received 5 sessions of paired tone pips (conditioned stimulus, CS) and foot shocks (unconditioned stimulus, US), while control mice received 5 sessions of just tone pips (Figure 1A). Mice were tested with the conditioned stimulus 24 hours after conditioning. All conditioned mice (*n* = 3) exhibited high freezing responses, while all control mice (*n* = 3) exhibited low freezing responses (Extended Data Figure 2). At 4 days after conditioning, we harvested auditory cortex (A1) tissue blocks from these mice and prepared them for ssEM (Figure 1B). We sectioned a total of 2.8 × 10^5^ μm^3^ A1 tissue at 50 nm section thickness, and imaged at 2-4 nm lateral resolution (Supp. Video 1).

**Figure 2.**
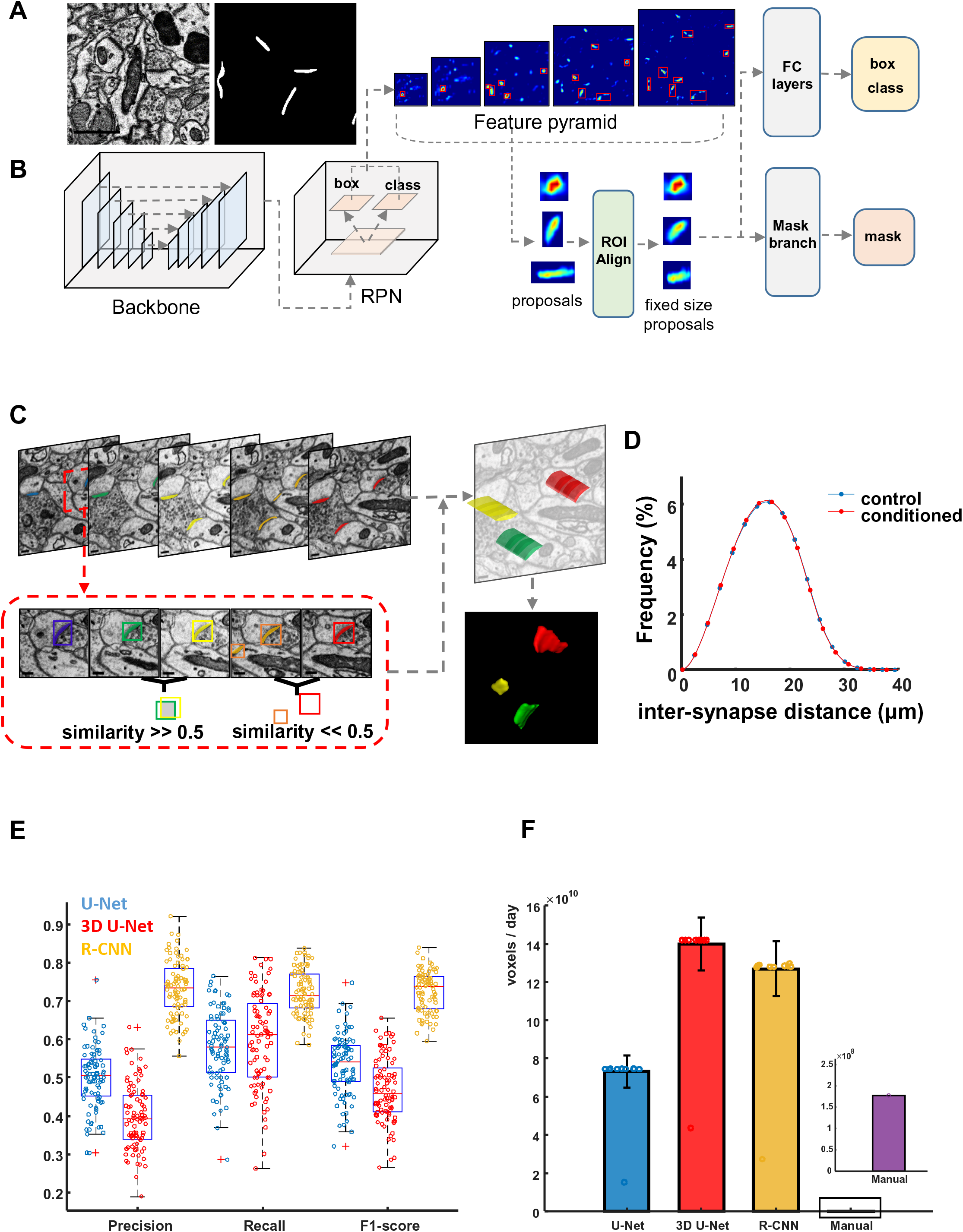
Automated identification and reconstruction of synapses. (A) Examples of an EM image and a binary synaptic mask predicted by Mask R-CNN. Scale bar: 1 μm. (B) Network architecture of Mask R-CNN, which takes a 1,024 × 1,024 image patch as input and outputs the bounding boxes and binary masks of all synapses in the input image (C) Sketch of the similarity-index-based 3D connection algorithm used for reconstructing synapses. The similarity of two synapses from adjacent layers depends on the intersection-over-union of the bounding boxes. (D) Normalized histograms of Euclidean distances between any two synapses in control (blue) and conditioned mice (red); note that these accord with a normal distribution (control: p = 0.1149, two-sided Kolmogorov-Smirnov test, conditioned: p = 0.4801, two-sided Kolmogorov-Smirnov test). (E) Comparison with other state-of-the-art methods in terms of precision, recall, and F1-score metrics on a public ssEM dataset. Each circle indicates an image from the test sets (half of this public ssEM dataset). (F) Comparison with other automatic state-of-the-art CNN methods as well as manual annotation on processing speed, which was here evaluated as the number of voxels processed per day. With one GPU, R-CNN (golden) is in the same order of magnitude as 3D U-Net (red), one order of magnitude faster than U-Net (blue), and nearly three orders of magnitude faster than manual processing by humans (purple).

### Deep learning-based reconstruction of synapses in A1

We first explored the synaptic changes associated with AFC, by extracting structural information of synapses from ssEM data. Synapses have distinct ultrastructural properties: pre- and postsynaptic membranes with a synaptic cleft in between, postsynaptic density (PSD) and abundant synaptic vesicles in presynaptic terminals. These special features enabled us to design a 2D-3D pipeline to detect and reconstruct synapses at the two-dimensional (2D) and 3D levels. Due to the high anisotropy of voxels (x-y resolution: 2-4 nm, z: 50 nm) and intrinsic local misalignments in most ssEM data, using a 3D convolutional neural network (3D CNN) increases the computational complexity without offering any improvement in performance. Therefore, we used the Mask R-CNN^18^ model at the 2D level to detect and segment synapses in each 2D image (Figure 2A). The Mask R-CNN is a deep neural network for the instance segmentation, which can separate distinct objects in an image. As illustrated in Figure 2B, Mask R-CNN is composed of three primary parts: backbone network, Region Proposal Network (RPN) and Region-CNN (R-CNN). The backbone network provides shared feature maps for the other two parts. The backbone used for synapse detection is a Feature Pyramid Network (FPN, Figure 2B, Supp. Methods)^19^, which we modified from the ResNet50 model^20^. As a typical two-stage detector, it first generates enough region proposals to guarantee the pre-specified recall rate with a RPN. Subsequently, the feature maps of the proposals are extracted as Regions of Interest (RoIs). R-CNN then makes further classification (predicting the scores being synapses or not), regression (predicting the coordinates of synapses’ bounding boxes) and predicts a pixel level mask for the RoIs identified in the first stage. The classification branch that predicted each RoI as a synaptic or non-synaptic object explored both the features of the synaptic vesicles and the PSD. This second stage guarantees the precision rate. After obtaining the final positions, the mask branch predicted the segmentation masks of the detected PSDs.

To train the network, a total of 600 ssEM images from the aforementioned mouse A1 dataset were annotated by two expert annotators with cross-validation (two volumes of 2,048 × 2,048 × 300 voxels for the control and conditioned groups, respectively), which was split into training (60%), validation (20%), or test (20%) sets. Evaluation against the test set showed that our pipeline achieved a 0.90 precision rate and a 0.83 recall rate for synapse detection (Extended Data Figure 3A and 3C).

**Figure 3.**
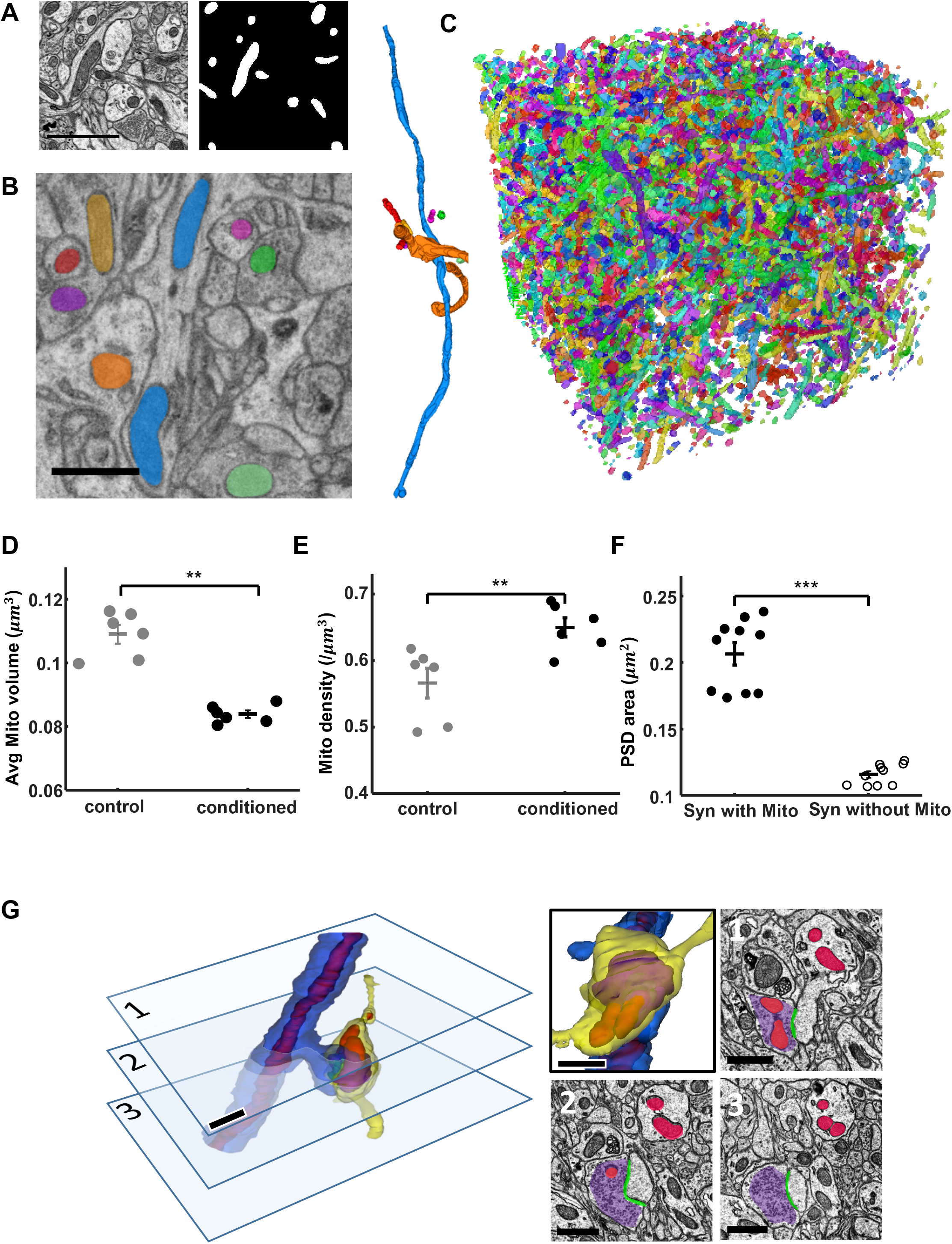
Automated identification of mitochondria and vesicle clouds. (A) Examples of an EM image and a binary mitochondrial mask predicted by Mask R-CNN. Scale bar: 1 μm. (B) EM image and 3D visualization of 8 mitochondria located in dendrites or axons. Scale bar: 1 μm. (C) 3D visualizations of all mitochondria from one 22 × 24 × 25 μm image stack. (D) The average mitochondrial volume is smaller in conditioned mice (0.08 ± 0.001 μm^3^) than in controls (0.11 ± 0.003 μm^3^). p < 0.01, two-sided t-test. (E) The mitochondrial density is higher in conditioned mice (0.65 ± 0.014 μm^3^) than in controls (0.57 ± 0.023 μm^3^). p < 0.01, two-sided t-test. (F) The postsynaptic density (PSD) area of synapses with presynaptic mitochondria (0.21 ± 0.008 μm^2^) is larger than the PSD area of synapses without presynaptic mitochondria (0.12 ± 0.002 μm^2^). p < 0.001, two-sided t-test. (G) A typical example of a spine synapse with the identified PSD (green), mitochondria (red) and vesicle cloud (purple). Another view and the corresponding EM images are displayed on the right. Scale bar: 1 μm.

To quantitatively evaluate the performance (efficiency and accuracy) of the Mask R-CNN, we compared it against other state-of-the-art CNNs using a previously reported public ssEM dataset^1^ comprising 178 slices sized 8576 × 7616 pixels. The dataset is divided into two equal parts, one for training and one for testing. The baseline network U-Net^21^ and the 3D U-Net^22^ are commonly used for biomedical image segmentation tasks. In terms of precision rate, recall rate, and F1-score (the harmonic mean of precision and recall), our pipeline outperformed U-Net and 3D U-Net (Figure 2E).

We also compared the processing speed of the three networks equipped with one graphics processing unit (GPU) as well as that of manual annotation (Figure 2F). The results confirmed that our method ( 1.27 × 10^11^ ± 1.44 × 10^10^ voxels/day) is comparable to 3D U-Net ( 1.40 × 10^11^ ± 1.39 × 10^10^ voxels/day), one order of magnitude faster than U-Net (7.33 × 10^10^ ± 8.40 × 10^9^ voxels/day), and almost three orders of magnitude faster than manual annotation (1.77 × 10^8^ voxels/day).

At the 3D level, we used a 3D connection algorithm (Figure 2C) to find the instance-level connected components and reconstruct synapses. Based on the continuity of the aligned ssEM volume and the spatial structure of the synapses, we constructed similarity matrices (Supp. Methods) between adjacent layers with synapse detection boxes. If the similarity of the two bounding boxes was greater than a certain threshold (default set at 0.5), we considered the corresponding synapse to be the same one at the 3D level. If a synapse appeared in 3 continuous layers or more, it was retained and given a unique label; otherwise, it was discarded as a false positive. Therefore, this connection algorithm was also used as a post-processing method to remove false positives and refine the segmentation and detection results. After connected component labeling, we could obtain the synaptic instance segmentation results where each label indicated a unique synapse in 3D.

Using our 2D-3D pipeline, we automatically identified, segmented, and reconstructed over 160,000 synapses from 12 A1 tissue blocks of 6 mice (Extended Data Figure 4, Supp. Video 2). To compare the spatial distribution of synapses in control and fear conditioned animals, we computed the distance between any two synapses, and found that synapses were uniformly distributed both in control and conditioned animals, with no difference in the average inter-spine distance (Figure 2D). Therefore, AFC does not cause a major change in synapse number or distribution in A1.

**Figure 4.**
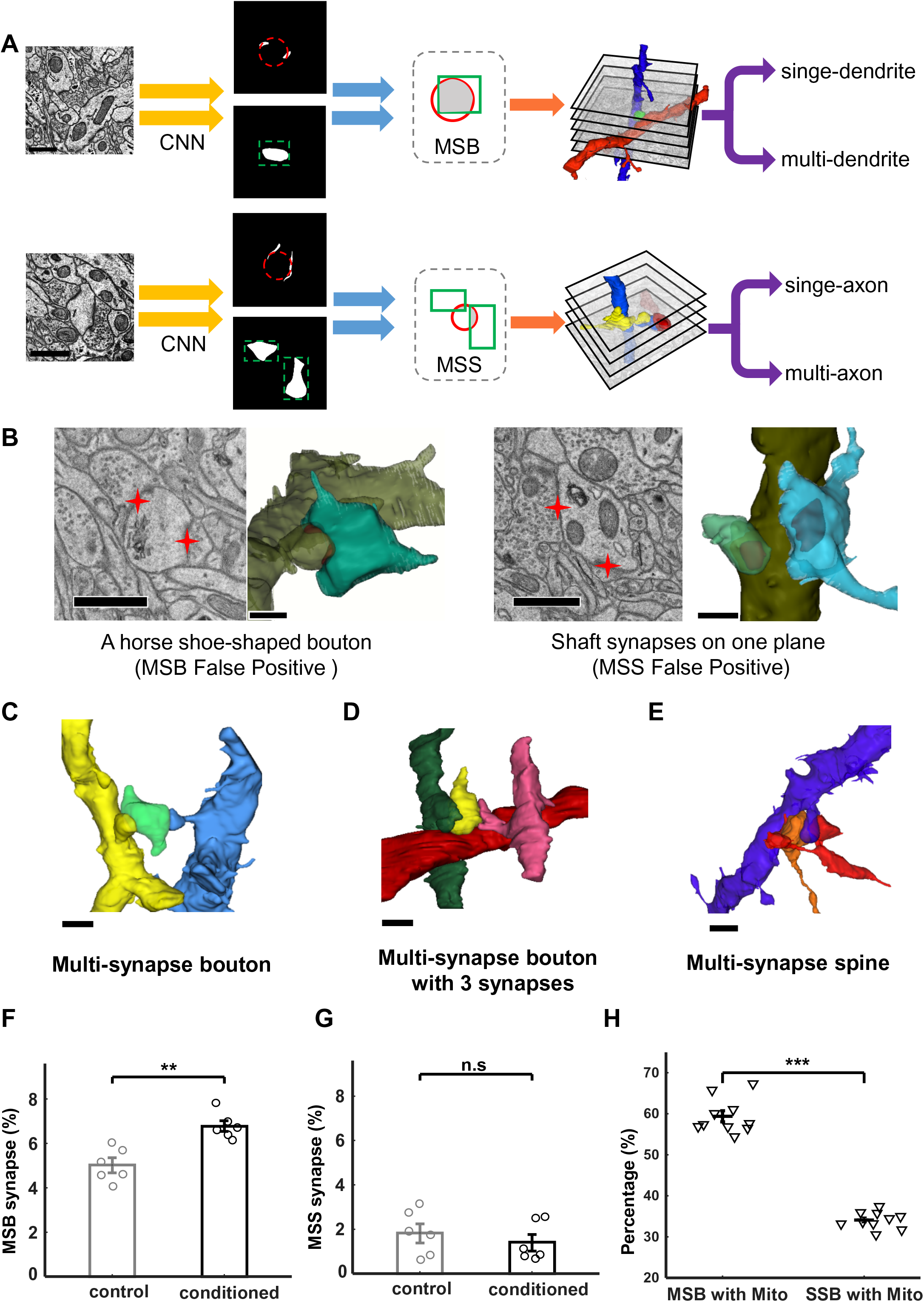
Synaptic organization in A1. (A) MCS detection and classification. MCS candidates are first automatically marked based on a close-vicinity criterion, and then manually verified and classified as MSBs or MSSs by estimating the number of boutons based on vesicle cloud segmentation. Specifically, the bouton number is estimated by intersecting the fitting circle of the synapse and the bounding box of the vesicle cloud. Neurite reconstructions are then introduced to determine the origination of multiple spines or boutons. Red circles: fitted circles by synapse segmentation; Green boxes: minimum boundary rectangles of vesicle cloud segmentation. Scale bar: 1 μm. (B) Two examples of false positives identified by 3D reconstruction of synapses. The left subpanel is an MSB false positive that was falsely detected owing to the horse shoe-shaped bouton (dark green). The right subpanel is an MSS false positive comprising two shaft synapses. Scale bar: 1 μm. (C-E) Examples of a 3D reconstructed MSB (C), an MSB with more than 2 postsynaptic sites (D), and an MSS (E). Scale bar: 1 μm. (F) The percentage of MSB synapses is higher in conditioned mice (6.8 ± 0.2 %) than in controls (5.0 ± 0.3 %). p < 0.01, two-sided t-test. (G) There is no difference in the percentage of MSS synapses between control (1.8 ± 0.4 %) and conditioned mice (1.4 ± 0.4 %). p = 0.46, two-sided t-test. (H) The percentage of MSB synapses with presynaptic mitochondria (59.41 ± 1.3%) is higher than the percentage of SSB synapses with presynaptic mitochondria (34.12 ± 0.6%). p < 0.001, two-sided t-test.

### Deep learning-based reconstruction of mitochondria in A1

We designed our 2D-3D pipeline such that it is capable of identifying any cellular compartment or organelle with borders and distinct structural properties. We then used this pipeline to identify mitochondria, the major energy source for cellular functions and neuronal activities, from the same ssEM dataset. Mask R-CNN first predicted a binary mitochondrial mask (Figure 3A) for each input image, after which the 3D connection algorithm produced the reconstructed mitochondria (Figure 3B). To build a ground truth mitochondria dataset, mitochondria from 20 images (7,492 × 7,492 pixels) were labeled by experienced annotators. The train-validation-test split ratio used here was the same as for synapses. To correct for discrepancies in imaging conditions, we preprocessed the images using histogram matching. The images were cropped into smaller patches (1,024 × 1,024 pixels) for training the R-CNN. The proposed algorithm achieved a 0.93 precision rate and a 0.91 recall rate for mitochondria detection on the test set (Extended Data Figure 5).

**Figure 5.**
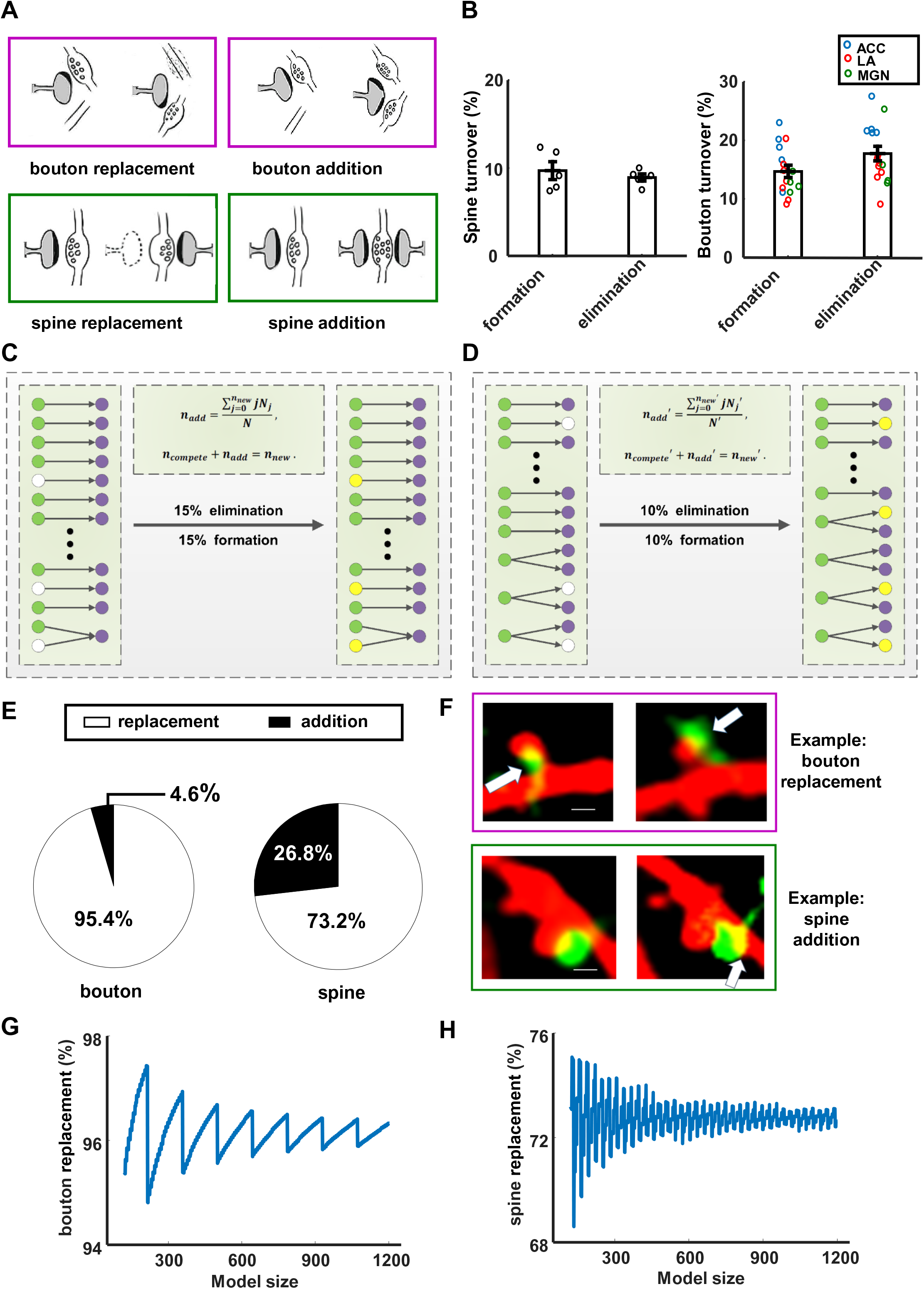
Synaptic turnover patterns. (A) Cartoons showing replacement and addition in bouton and spine turnover. (B) Four-day turnover rates of spines (formation: 9.71 ± 1.02 %, elimination: 8.93 ± 0.41 %) and boutons (formation:14.73 ± 1.03 %, elimination: 17.76 ± 1.24 %) in the auditory cortex of fear conditioned mice. Each circle represents data from a single mouse. The bouton turnover rate is averaged among axons that project to the auditory cortex from the lateral amygdala (LA), the anterior cingulate cortex (ACC), and the medial geniculate nucleus (MGN). (C-D) Diagrams showing bouton (C) and spine (D) turnover in combinatorial mathematical models. Green: existing boutons. Purple: exisiting spines. White: eliminated spine/bouton; yellow: added new spine/bouton; arrow: synaptic connection. (E) Estimation of the percentage of replacement and addition for boutons (left) and spines (right) based on combinatorial mathematical modeling. Models predict 95.4% replacement of boutons and 73.2% replacement of spines. (F) Synaptic turnover examples from *in vivo* two-photon microscopy analysis. Arrows point to elimination of an old bouton and to formation of a new bouton (green, top), and to an addition of a new spine (red, bottom). Scale bar: 1 μm. (G) The percentage of bouton replacement is around 96% when the model size is scaled up to 12,000 synapses. (H) The percentage of spine replacement is around 72% when model size is scaled up to 12,000 synapses.

After validation against the ground truth dataset, we applied the 2D-3D pipeline to examine mitochondrial changes after AFC. We identified, segmented, and reconstructed over 135,000 mitochondria from 12 A1 tissue blocks of 6 mice (Figure 3C, Supp. Video 3). By combining the synapse dataset and mitochondria dataset, we found that synapses with presynaptic mitochondria had larger PSDs than those without (PSD size with presynaptic mitochondria: 0.21 ± 0.008 μm^2^, PSD size without: 0.12 ± 0.002 μm^2^, Figure 3F). AFC significantly decreased mitochondrial volumes (control: 0.11 ± 0.003 μm^3^, conditioned: 0.08 ± 0.001 μm^3^, Figure 3D) while increased mitochondrial density (control: 0.57 ± 0.023 μm^3^, conditioned: 0.65 ± 0.014 μm^3^, Figure 3E). Thus although AFC does not alter the spatial distribution of synapses, it increases the number and decreases the size of mitochondria, suggesting that mitochondrial fission may accompany AFC.

### A synapse dataset combined with mitochondria and synaptic vesicles

We successfully identified and reconstructed the PSDs of synapses from ssEM images with our 2D-3D pipeline, but the direction of the synaptic connectivity is still unclear. Vesicle clouds in the presynaptic terminals provide essential information for distinguishing axons from dendrites in ssEM images. Thus, we applied FusionNet^23^, a variant of U-Net, to detect the synaptic vesicle clouds (Extended Data Figure 6A). FusionNet used the residual blocks and summation-based skip connections (Extended Data Figure 6C), which could achieve state-of-the-art performance in segmenting the ssEM data. The network predicted probability maps each element indicating the probability of belonging to the foreground. Synaptic vesicles were about 60 nm in diameter and concentrated in the presynaptic region. Since labeling each synaptic vesicle is extremely time-consuming, we annotated vesicle clouds to reduce the annotation workload. We extracted two volumes (2,048 × 2,048 × 50 voxels) for synaptic vesicle annotation, and divided them into training, validation, and test sets using the same split ratio as for synapses and mitochondria. To evaluate the performance on the test set, a thresholding operation was conducted, and FusionNet yielded a precision rate of 0.83 and a recall rate of 0.80 (Extended Data Figure 6B).

**Figure 6.**
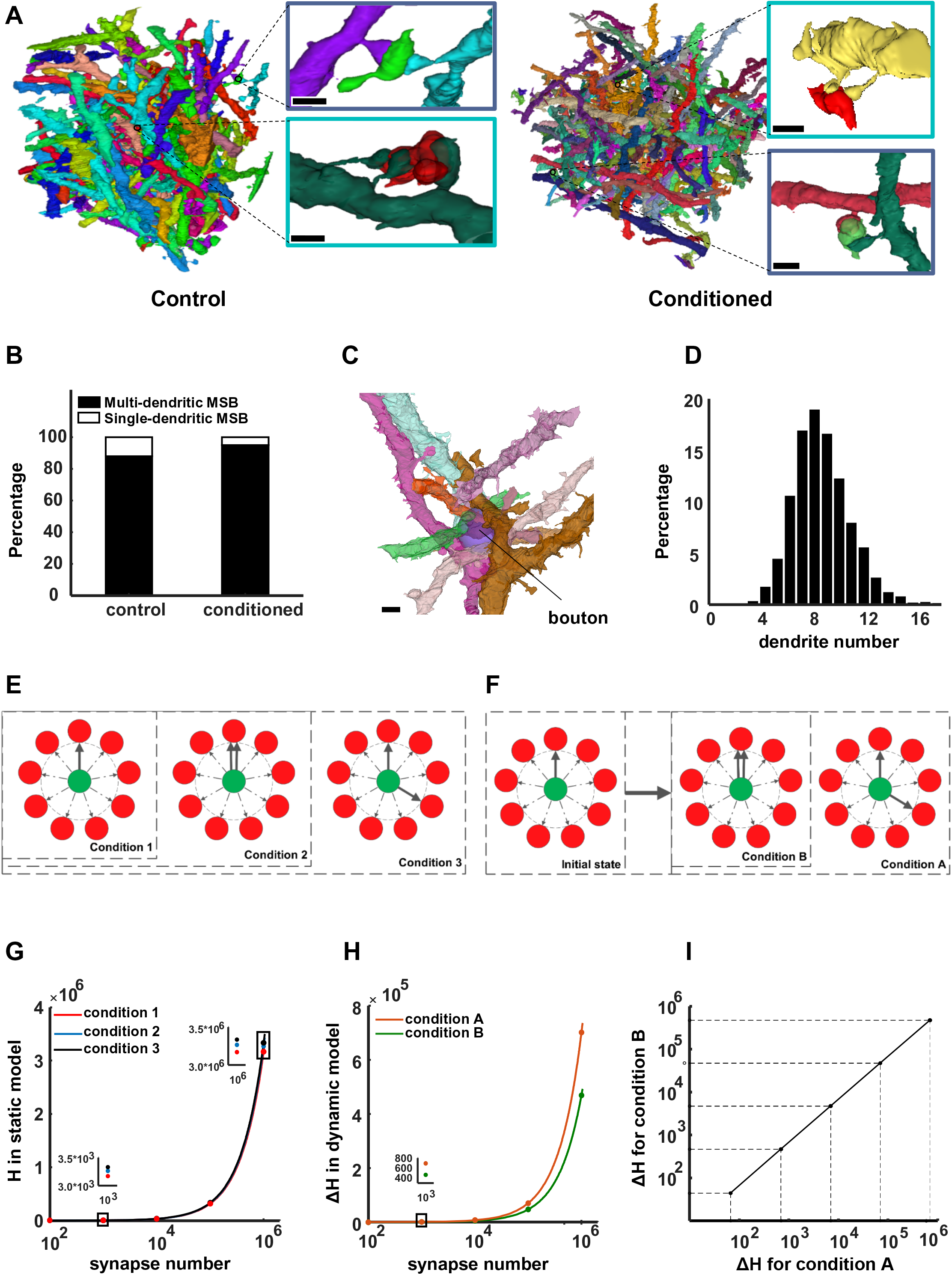
Saturated reconstruction and information storage capacity calculation. (A) Two reconstructed tissue blocks from control (left: 22 × 24 × 25 μm) and conditioned mice (right: 33 × 28 × 25 μm). Insets show examples of single- or multi-dendritic MSBs. Scale bar: 1 μm. (B) Percentage of single- and multi-dendritic MSBs in control (multi-dendrite MSBs: 86%) and conditioned mice (multi-dendrite MSBs: 95%). (C) A reconstructed example showing 10 dendrites intersecting within a 1-μm radius of a bouton. Scale bar: 1 μm. (D) Distribution of the number of dendrites within a 1-μm radius of a bouton. n = 5774 boutons. (E) Illustrations showing synaptic connectivity patterns of a static model. Condition 1: 1-to-1 synapses only; condition 2: 1-to-1 synapses and single-dendritic MSBs; condition 3: 1-to-1 synapses, single- and multi-dendritic MSBs. (F) Illustrations showing synaptic connectivity patterns of a plastic model. Condition A: multi-dendritic MSBs, condition B: single-dendritic MSB only. Red: dendrite; green: bouton; solid arrow: one possible synaptic connection; dashed arrow: potential synaptic connection. (G) Information storage capacity (ISC) for static networks of different sizes for 3 conditions: 1-to-1 synapses only (condition 1); 1-to-1 synapses and single-dendritic MSBs (condition 2); 1-to-1 synapses, single- and multi-dendritic MSBs (condition 3). (H) Increase in ISC for plastic networks of different sizes for 2 conditions: new synapse on any passing dendrite (condition A); new synapse on the same dendrite (condition B). (I) Linear relationship for the increase in ISC for conditions A and B across different network sizes.

After applying the trained version of FusionNet to identify vesicle clouds in A1, we constructed a large-scale dataset containing synapses, mitochondria and vesicles from control and conditioned mice, which can be used then for studying the cellular ultrastructural changes associated with AFC. An example showing synaptic ultrastructure, including synaptic cleft, mitochondria, and vesicle cloud, is presented in Figure 3G.

### Fear conditioning increased a specific type of multi-contact synapses

Synapses that form 1-to-N or N-to-1 connections, termed multiple-contact synapses (MCSs), have been observed in the brains of mice, rabbits and monkeys^24–26^, and were implicated in functions such as eye-blink conditioning^27^. But statistical and structural analyses were limited due to the small sample size obtained by manual notation of EM data in previous studies. In our previous work, we found that in adult mouse A1, synaptogenesis rarely occurs *de novo*, but rather by addition of new boutons or spines to existing synapses, termed “partial addition rule”^3^. The finding that AFC leads to an increase in spine and bouton formation together with this rule may lead to an increase in MCSs, if the additions are not accompanied by eliminations of existing synapses via synaptic replacement.

To find out if MCSs serve as a synaptic memory engram for AFC, we designed a semi-automated method to localize all MCSs in the tissue blocks to examine MCSs in a large scale (Figure 4A and Extended Data Figure 7A). To identify MCSs from annotated synapses, we took advantage of the fact that MCSs share either a bouton or a PSD, so the distances of PSDs or boutons of one MCS are restricted to the size of a synapse (∼1-µm). Based on this biological prior information, the candidate MCSs were detected by restricting the distances between identified synapses. Then expert annotators proofread to exclude the false positives (Figure 4B) which were not easily distinguished in 2D images. By combining the segmentation results of synapses and vesicle clouds to estimate the number of boutons in one MCS, we then classified all the identified MCSs into two types: those consisting of a single bouton contacting multiple PSDs (“Multi-Synaptic Bouton”, MSB, Figure 4C and 4D) or those consisting of a single spine contacting multiple boutons (“Multi-Synaptic Spine”, MSS, Figure 4E).

We found that the percentage of MSB synapses was significantly increased after AFC (control: 5.0 ± 0.3 %, conditioned: 6.8 ± 0.2 %, Figure 4F), whereas that of MSS synapses did not change (control: 1.8 ± 0.4 %, conditioned: 1.4 ± 0.4 %, Figure 4G). We also noted that the majority (∼98.58 ± 0.4%) of MSBs had 2 postsynaptic targets, yet a few had 3 or more (Figure 4D), and the percentage of MSBs forming more than 2 synapses was significantly elevated in the conditioned animals (control: 1.07 ± 0.3%, conditioned: 2.26 ± 0.3 %, Extended Data Figure 7D). Together, these results showed that AFC affected MCSs, specifically by promoting formation of MSBs. We also found that the percentage of MSB synapses with presynaptic mitochondria is significantly higher than that of single-synaptic bouton (SSB) synapses with presynaptic mitochondria (MSB: 59.41 ± 1.3%, SSB: 34.12 ± 0.6%, Figure 4H), suggesting that MSBs are more energy-demanding.

### Combinatorial modeling to assess bouton and spine turnover patterns

Synapses undergo constant turnover in the adult brain, and long-term memory storage may involve the formation of new synapses^1–3^. However, it is unclear to what extent such synaptic additions may be accompanied by the elimination of existing synapses (or whether they tend to co-exist). We performed *in vivo* two-photon imaging of fluorescently labeled boutons and spines in A1 of control and fear conditioned mice, and computed the bouton and spine turnover rates at 4 days after AFC. The formation and elimination rates of spines over the 4-day period are 9.7 ± 1.02% and 8.9 ± 0.41%, and of boutons 14.7 ± 1.03% and 17.8 ± 1.24%, with no significant net increase or decrease. We reasoned that for any synapse, if the addition of a new bouton/spine is always accompanied by the elimination of the existing bouton/spine (Figure 5A, left), then the percentage of MCSs should remain constant, otherwise there should be an increase of MCSs (Figure 5A, right). Thus, our finding that the percentage of MSSs remained unchanged after AFC (Figure 4G) suggests that new boutons tend to replace old ones, representing old connectivity patterns supplanted by new ones. In contrast, the significant increase in the number of MSBs (Figure 4F) suggests that new spines are added to existing synapses without eliminating old ones, representing addition of new connections while preserving old ones.

To quantitatively assess the difference in the turnover pattern of boutons and spines, specifically, to estimate the proportions of newly formed boutons/spines being replaced and/or added, we developed a combinatorial mathematical model that exhausted all turnover possibilities based on results from ssEM and *in vivo* imaging (Figure 5C and 5D). We used the model to estimate: 1) the proportion of boutons/spines for which formation was accompanied by elimination, and 2) the proportion of boutons/spines that are simply added to existing synapses without eliminating old ones.

We took the MCS/1-to-1 synapse composition of control mice as the starting situation and that of conditioned mice as the end situation, using a bipartite graph to model synaptic connections. In order to better reflect the difference in the MSB ratio before and after learning, as well as to take into account computational complexity, we modeled using 120 synapses to obtain the final expected values, using combinatorics to calculate the possibility of different patterns. We assumed an equal possibility for all turnover patterns. The percentages of MSBs and MSSs, and the elimination and formation rates of spines and boutons were all based on experimental data (Figure 4F, 4G and 5B). Synaptic turnover patterns that involve newly formed boutons/spines include the following 6 categories (‘-’ represents elimination and ‘+’ formation):

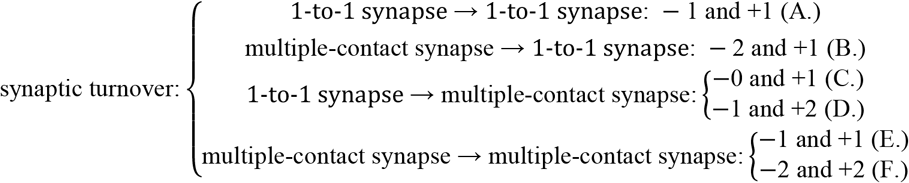

According to the above classification, “synaptic replacement” includes (A.), (B.), (D.), (E.) and (F.), and “synaptic addition” includes only (C.). We built two models, one for bouton turnover and MSS, the other for spine turnover and MSB. These two models capture bouton and spine turnover patterns, respectively (for detailed calculation, see Supp. Methods).

Our combinatorial mathematical modeling predicts a much higher percentage of replacement of boutons than that of spines (Figure 5E, 5G and 5H, Supp. Methods), which is consistent with our observation *in vivo*: among 856 putative synapses identified from two-photon imaging of the A1, there were 9 cases of a new bouton replacing an old bouton, and 4 cases of a new spine adding onto an existing synapse; there were no cases wherein a new bouton was added to an existing synapse (Figure 5F). Note that we attribute the low frequency of turnover events of MCSs present in the *in vivo* dataset to the sparseness of neuronal labeling. Together, these data suggest that each bouton tends to be the sole input of its postsynaptic counterpart, whereas spines can co-exist on a single bouton, suggesting the information transfer pattern of “1-to-N” is more common than “N-to-1” in synaptic transmissions.

### Evaluating the information storage capacity of MSBs in static synaptic networks

Compared to two 1-to-1 synapses, two synapses formed by one MSB saved cellular resources required for two distinct presynaptic boutons. To further investigate the information coding capacity of MSBs, we developed a mathematical model to calculate the information storage capacity (ISC)^28–31^ of a synaptic network comprising a set number of synaptic connections. The ISC of a synaptic network is given by the Shannon’s information entropy^32^ *H*, which measures the average uncertainty in the synaptic connection patterns of the network, and can be expressed as:

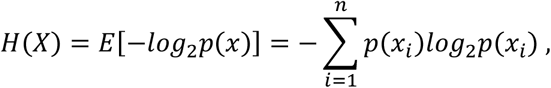

where, *n* is the number of all possible connection patterns, *X* is a random variable of synaptic connection patterns, *p*(*x*) is the probability mass function of *X*, and *p*(*x*_*i*_) is the probability measure of the occurrence of the i^th^ synaptic connection pattern *x*_*i*_. We assume that the probability of each pattern of synaptic connection is equal, and the model transforms into a mode to solve for the number of eligible patterns. Here, the degree of uncertainty in the network depends on the number of available dendrites to which each bouton could connect to. For cases where the multiple postsynaptic structures of one MSB originate from the same dendrite, we consider the connection to be the same as a 1-to-1 synapse. Therefore, the ISC of an MSB synapse is determined by how many dendrites that the MSB can connect to.

To determine the number of available dendrites for each bouton and the percentages of both single-dendritic and multi-dendritic MSBs, we used the Multicut pipeline^33^ and performed saturated reconstruction for two A1 tissue blocks of one control (22 × 24 × 25 μm) and one fear conditioned (33 × 28 × 25 μm) mouse (Figure 6A, Supp. Video 4-7, Extended Data Figure 8, Supp. Methods). Since the typical length of a spine^34^ is 1-µm, we calculated the number of dendrites passing the vicinity of a bouton within a 1-µm radius in the reconstructed data set (Figure 6C, Supp. Video 8, Supp. Methods). Among the 5,774 boutons analyzed (Figure S10), the median number of potential postsynaptic dendrites was 9 (Figure 6D), indicating that each bouton can potentially make synapses with 9 dendrites. We also traced each spine to its original dendrite to support categorization of MSB subtypes (single vs. multi-dendritic). Contrary to previous *in vitro* results reporting that LTP can lead to single-dendritic MSB—and thereby generating stronger connections between pre- and post-synaptic neurons^35^—we found that more than 90% of MSBs were connected to multiple dendrites (Figure 6B), forming 1-to-N connections.

We then calculated the ISC of the synaptic network based on the synapse reconstruction results containing the following types of connections: 1) 1-to-1 synapses only; 2) 1-to-1 synapses and single-dendritic MSBs; 3) 1-to-1 synapses, single- and multi-dendritic MSBs (Figure 6E). In a model with 100 synapses, of which 6% are MSB synapses (based on data from Figure 4F), single-dendritic MSB increased ISC by 2.5% over 1-to-1 synapses, and multi-dendritic MSB further increased 2.2%. The benefits of adding MSBs remained when the model was scaled up to 10^6^ synapses (Figure 6G, Supp. Methods). These results indicate that MSBs in a static network do increase information coding capacity, but only slightly.

### A plastic connectivity model for the information storage capacity of synaptic networks

The percentage of multi-dendritic MSBs among all MSBs was higher in the conditioned mouse (95.1%) than in the control mouse (87.0%). This result, together with the percentages of MSBs in control and conditioned mice (5.0 ± 0.3% vs. 6.8 ± 0.2%, Figure 4F), indicates that essentially all of the MSBs newly formed after AFC were multi-dendritic. Thus, AFC resulted in boutons making novel connections with other dendrites, rather than strengthening existing connections.

To evaluate the difference in ISC from establishment of new synaptic connections under the two conditions, multi-dendritic vs. single-dendritic, we built a synaptic network model that incorporated synaptic plasticity by adding 10% more spines to the boutons (Figure 6F), based on the *in vivo* imaging results (Figure 5B). The increase in ISC depends on the number of new connections formed, and the number of potential connection targets for each new connection. For newly formed multi-dendritic connections, the number of potential targets for each new connection is the number of dendrites that a bouton can make contact with (Figure 6D; condition A, forming multi-dendritic MSB); for single-dendritic, the number of potential target is 1 (condition B, forming single-dendritic MSB).

The changes in ISC due to this 10% addition of synapses in the network, denoted by the increase of information entropy Δ*H*, were calculated for the condition A and condition B as follows:

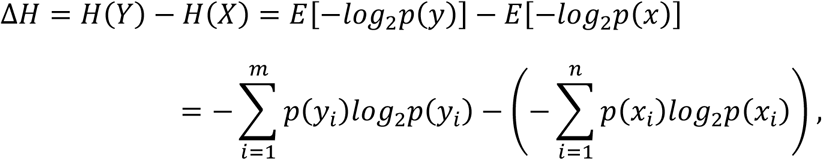

where, *n* and m are the number of all synaptic connection patterns before and after formation, respectively; *Y* and *X* are the random variables of the connection patterns after and before formation, respectively; *p*(*y*) and *p*(*x*) are the probability mass functions of *Y* and *X*, respectively; *p*(*y*_*i*_) is the probability of occurrence of the ith synaptic connection pattern after formation *y*_*i*_; *p*(*x*_*i*_) is the probability of the ith connection pattern before formation *x*_*i*_ . We assume that the probability of each possible connection pattern is equal, the above dynamic model can also transform into a mode to solve for the number of eligible patterns.

Notably, whereas a multi-dendritic MSB only slightly adds to the ISC in a static network (Figure 6G), in this plastic network, the increase in ISC for condition A was more than 50% higher (Figure 6H) than that of condition B. And we found that this relative advantage for multi-dendritic connectivity scaled linearly with network size (Figure 6I). This result indicates that establishing new connections in a plastic network by forming multi-dendritic MSBs can increase information coding capacity significantly, by 50%.

## Discussion

It is well-established that learning can induce neuronal plasticity in the brain. In previous *in vivo* imaging studies, only a limited population of synapses were investigated, and it is not known how learning changes the shape and size of mitochondria in the brain. In this study, we used deep learning techniques to automatically detect synapses and mitochondria in ssEM images. This method allowed us to examine hundreds of thousands of synapses and mitochondria, and our results demonstrated that synaptic and mitochondrial organization were significantly affected by the learning process. With a particular focus on synapses consisting of multiple synaptic elements, we were able to identify a specific form of MCSs consisting of a single bouton and multiple spines from different dendrites, which were indicated by mathematical modeling to confer higher information storage than single-contact synapses.

In this paper, we proposed a novel 2D-3D pipeline to outline the 3D morphology of synapses from ssEM images. The first step focused on detecting and segmenting instances on 2D slices with advanced R-CNN, and the second converted the 2D instances into 3D individuals. Compared with other CNN techniques, the experimental results demonstrated that our method greatly improved detection and recall performance. For the detection of MCSs, we first located them based on close-proximity criterion, then classified them into MSBs and MSSs using presynaptic vesicle information, which is another important feature for synapse identification^10^. We further obtained the dense reconstruction results by solving the graph partitioning problem, so that the multiple spines (or boutons) of MSBs (or MSSs) can be traced back to their original dendrites (or axons). This is the first report of detecting and classifying MCSs based on their connectivity. However, the inherent error of our MCS detection algorithm is a limitation of this study, for which predicting the synaptic partner neurons with CNN may be a solution in the future.

Dendritic spines have been extensively studied by *in vivo* imaging methods, because their unique shape makes them easily identifiable. However, there are also synapses formed on dendritic shafts, which are not visible by fluorescent imaging. They can, however, be easily identified using our synapse classifier algorithms (Extended Data Figure 9A, 9B, 9C and 9D) from ssEM images. In fact, 26% of all identified synapses were shaft synapses (Extended Data Figure 9G), and after learning, their number decreased as spine synapses increased (Extended Data Figure 9E and 9F), ultimately resulting in an unaltered total number of synapses. These synapses likely contribute to the homeostasis of cellular resource relocation and synapse organization.

The ssEM method enabled us to identify a special type of synapse, namely the multiple-contact synapses. Our previous work^3^ indicated that most new synapses are formed by adding a synaptic element, either a presynaptic bouton or postsynaptic spine, to an existing synapse. We called this the “partial addition rule”. Adding a bouton to an existing synapse creates an MSS, while adding a spine creates an MSB. We found that the MSB number increased significantly after learning, but MSS stayed the same. Since bouton formation and spine formation both increased after learning, this result suggested that a new bouton was more likely to compete and oust the previous bouton in a synapse for being the sole input to a postsynaptic spine, while the new spine and the old spine could co-exist on a single bouton, leading to a higher proportion of MSBs. MSS synapses may remain in a transient state during the switch from an old synaptic contact to a new one, while MSB represents a more stable synaptic configuration, allowing simultaneous information transfer from a single bouton to multiple postsynaptic sites. For any MSS, boutons from different neurons likely have unsynchronized activities, and only one of them would correlate best with the MSS. According to the Hebbian postulate “fire together, wire together”, only the best correlated bouton will win the competition to be the sole input for the spine, and the MSS will turn into a single-synaptic spine.

In most cases, the transmitters released from presynaptic sites outnumber the postsynaptic receptors, so it is efficient to transmit information from one bouton to multiple postsynaptic targets. Indeed, most MSBs made contact with more than one dendritic branch, broadcasting information from one neuron to multiple neurons with one multiple-contact bouton, with minimal cellular energy consumed. We noted that the proportion of MSB is relatively low, consistent with the notion that memory encoding may be very sparse. For any piece of memory, only very few synapses are involved in coding the information. Our model indicated a 50% increase in information storage capacity for multi-dendritic MSBs, which provides great potential for synaptic plasticity with minimal increase of synapse number and structures.

By design, our 2D-3D pipeline can be used to identify any cellular compartment or organelle with distinct structural properties, and we utilized it to identify mitochondria. Mitochondria adapt to the cellular energy requirements by highly dynamic fusion and fission^36^. Mitochondrial dynamics are also found to be related to synaptic transmission and plasticity. There was evidence to suggest that increasing mitochondrial fragments can promote synapse formation^37^. Our results showed significant decrease in size and increase in number of mitochondria along with changes of synaptic organization following fear conditioning, suggesting that mitochondria may play a role in learning by balanced fusion and fission.

The recently reported connectome of 1mm^3^ human cerebral cortex^11^ shows the substantial improvement of EM imaging speed and the powerful ability of deep learning. The biggest difference is that rather than performing large-scale saturated reconstruction, we conducted specific and local classification of synapses and mitochondria, which allows ultra-structural analyses of specific cellular organelles without resource-demanding heavy computation. As our method applies region-based CNN to identify objects, it can potentially be used to extract other discrete, distinct structures from ssEM data, such as Golgi apparatus and nucleus, making it a versatile tool for ssEM image processing.

## Acknowledgments

We thank Drs. Mu-ming Poo, Ju Lu, Yi Zuo and Margaret S. Ho for critical comments and suggestions. We also thank Linlin Li, Lina Zhang, Jingbin Yuan and Jinyue Guo for technical support; Jie Luo, Jiazheng Liu, Yi Jiang and Lu Wang for manual proofreading. This work was supported by grants from the Ministry of Science and Technology of China (2018YFC1005004) and the Natural Science Foundation of China (31970960) to Y.Y., the Strategic Priority Research Program of CAS (XDB32030200), Bureau of International Cooperation, CAS (153D31KYSB20170059) to H.H., and the Natural Science Foundation of China (61673381) to Q-w. X.

## Author contribution

H.H., Q.X. and Y.Y. conceived and guided the project. J.Q., Y.Y. and Y.K. performed the animal experiments and provided the samples. H.M. conducted EM imaging. G.L. implemented automated location and navigation system. X.C. performed alignment. J.L. and B.H. developed the algorithms. Z.L. established the mathematical models. L.S. performed visualization. Y.Y., J.L., J.Q. and Z.L. wrote the manuscript with contributions from all authors.

## Extended Data Figure legends

**Extended Data Figure 1.**
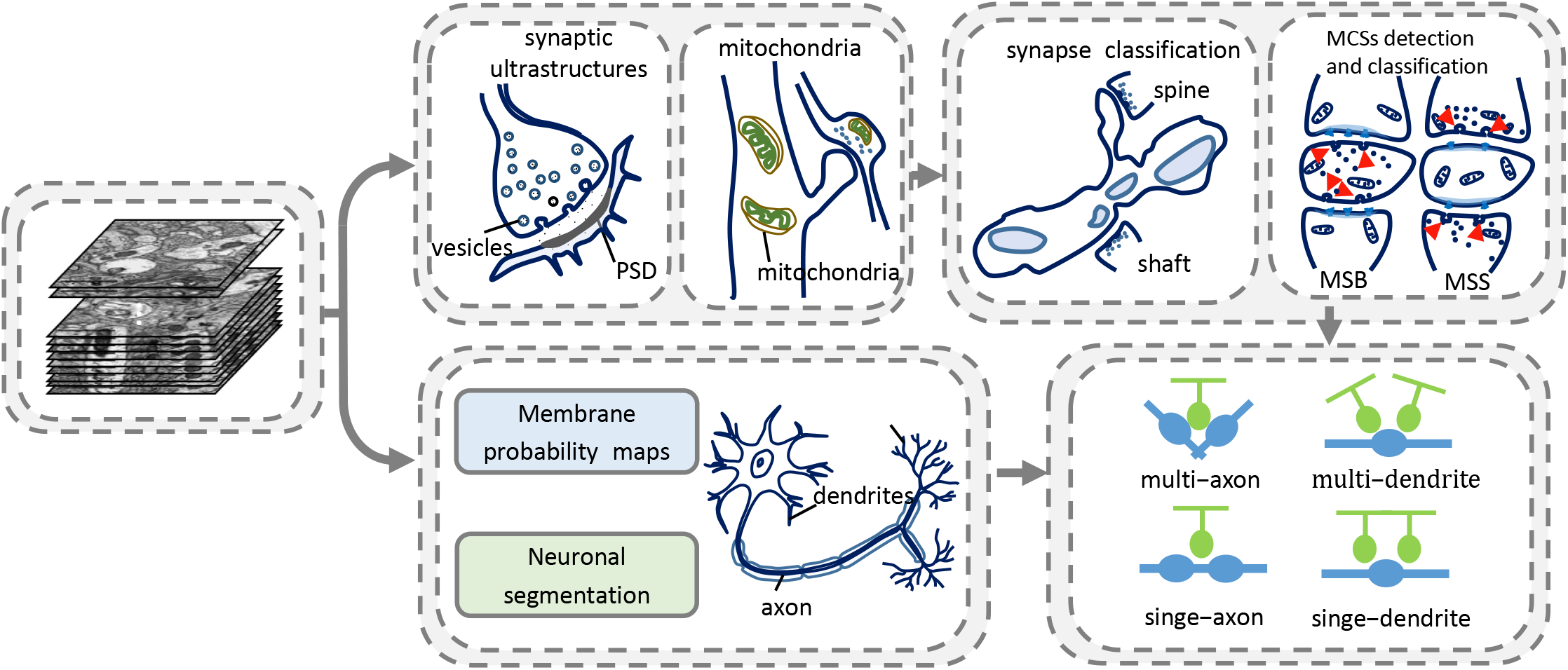
EM data processing procedure. The aligned EM stacks were first fed to R-CNNs for detection of the synaptic structures (PSD and synaptic vesicles), mitochondria, and neuronal membranes. The synapses were then classified into spine or shaft synapses based on the ultrastructural features. MCSs were identified and classified into MSBs and MSSs. Dense segmentation was achieved using the Multicut pipeline for determining the origination of the MCSs.

**Extended Data Figure 2.**
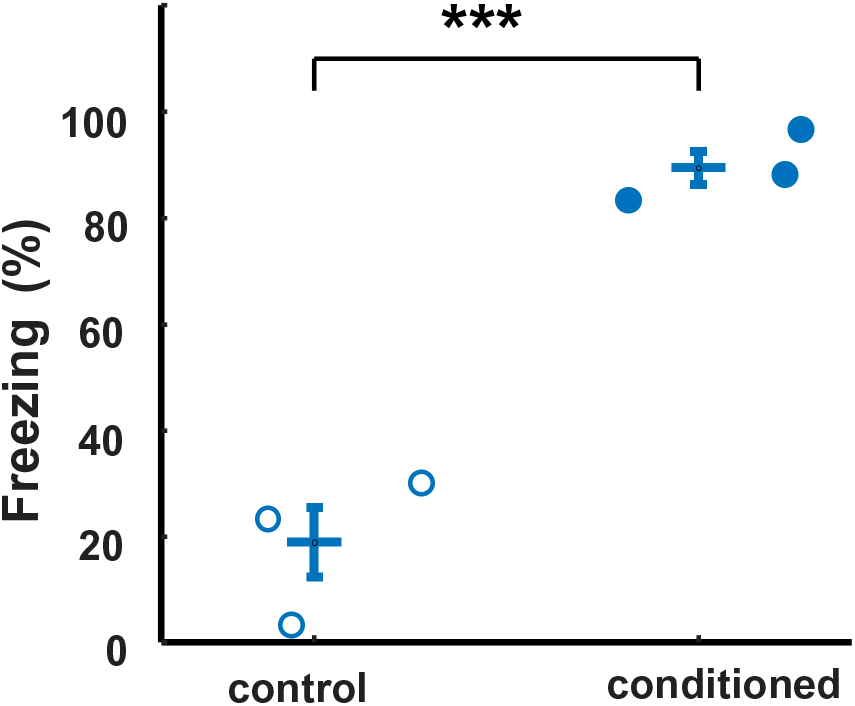
Behavioral results for auditory fear conditioning. Freezing levels of control (18.89 ± 8.01%) and fear conditioned (89.44 ± 3.89%) mice. Each circle represents data from one mouse. p < 0.005, Student’s t-test.

**Extended Data Figure 3.**
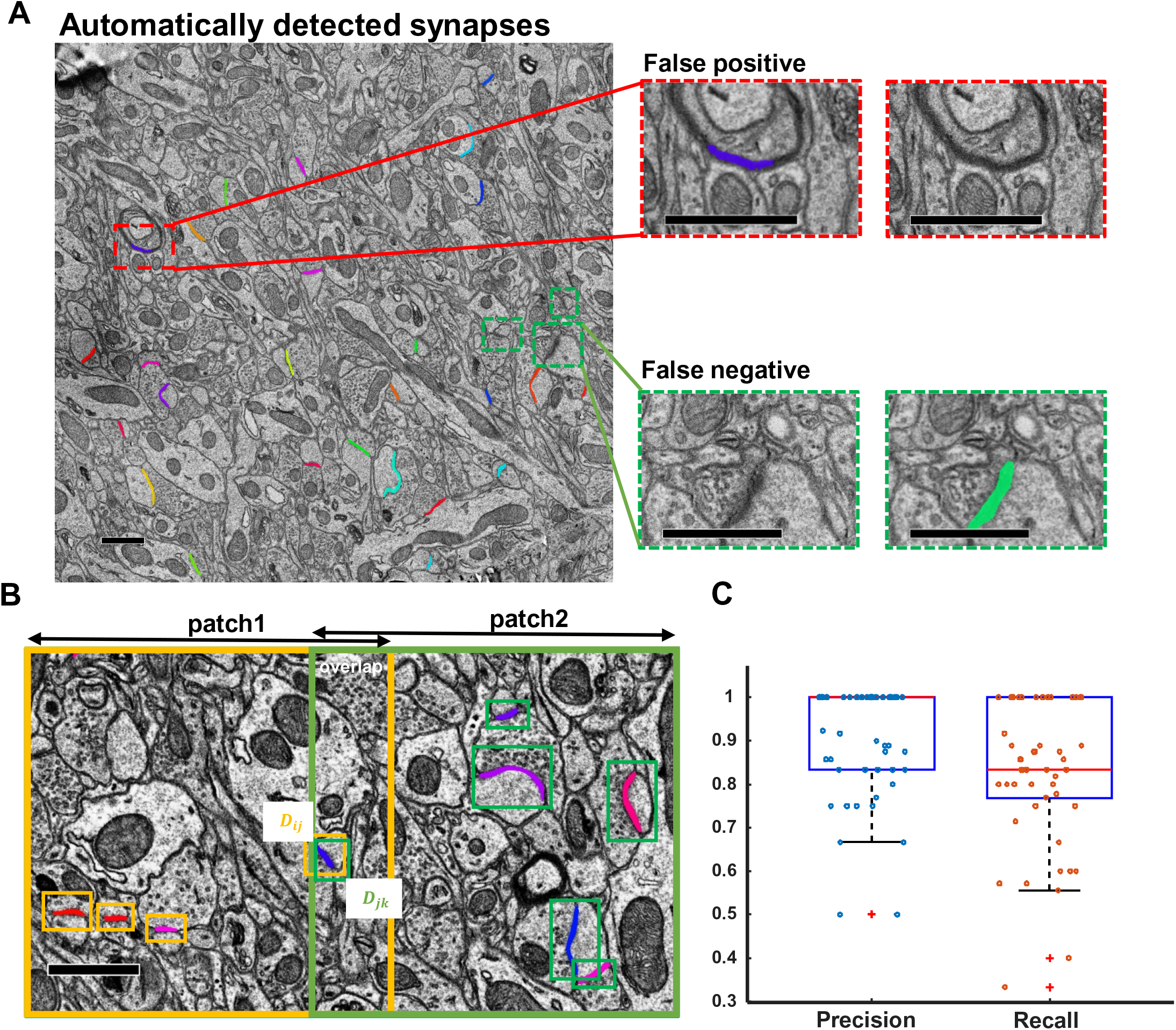
Automatic detection of synapses. (A) Examples showing automatically detected synapses in an EM image. Colored lines represent automatically detected synaptic clefts. Insets show false positives and false negatives by manual verification. Scale bar: 1 μm. (B) Diagram of single-layer fusion algorithm. Orange and green boxes represent two overlapping cropped patches. *D*_*ij*_ and *D*_*jk*_ reprenset repeated detection boxes in the overlapping region. Scale bar: 1 μm. (C) Evaluation of the synapse test set in terms of precision and recall. Each circle indicates one EM image from the test set.

**Extended Data Figure 4.**
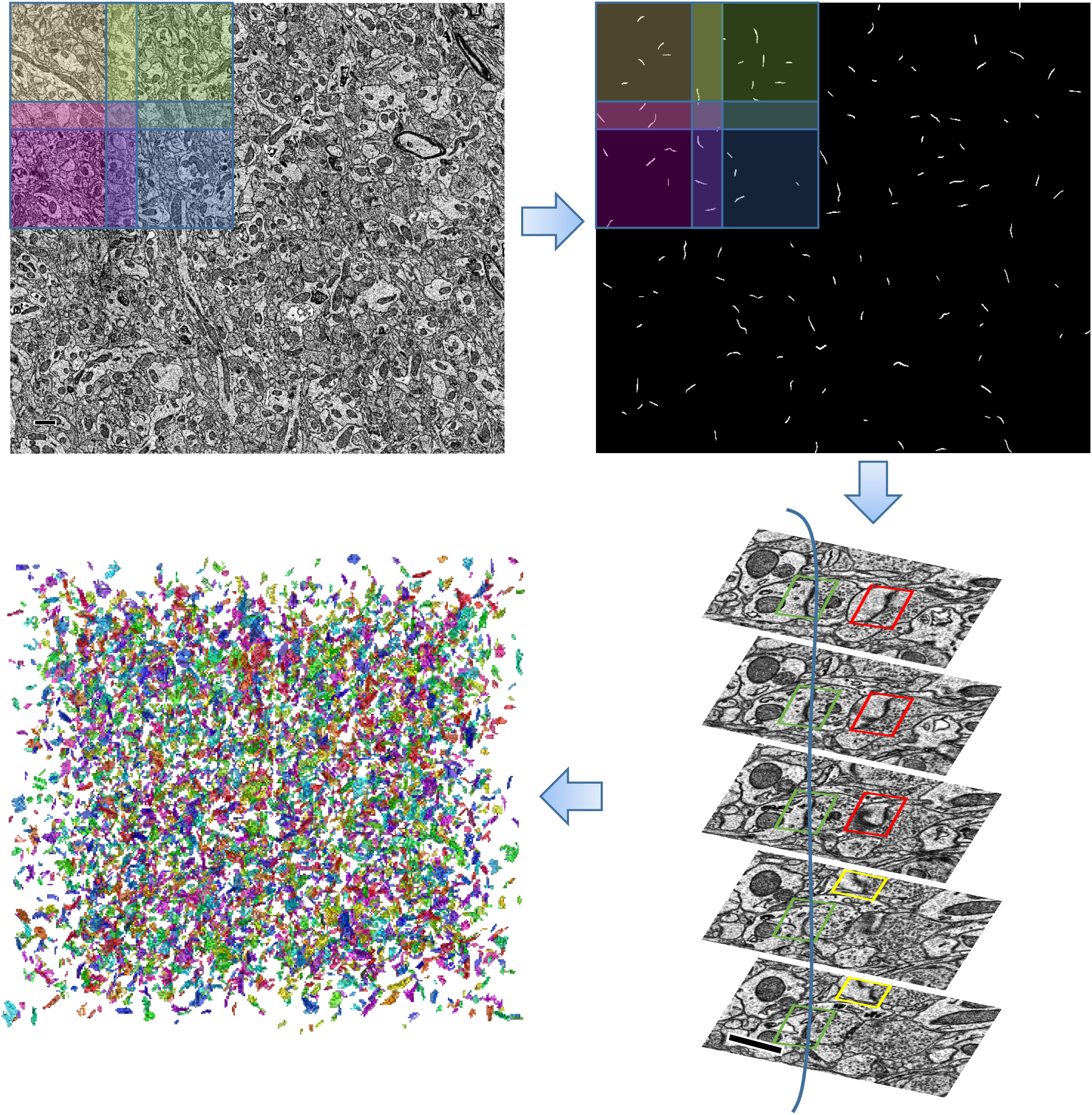
Pipeline for synapse reconstruction from ssEM images. The original EM images were cropped into overlapping small patches. The patches were then fed into the trained network for synapse identification, and the overlapping areas were fused using the single-layer fusion algorithm. The multilayer 3D connection algorithm was applied to obtain the reconstructed synapses, and 3D visualization was shown using ImageJ.

**Extended Data Figure 5.**
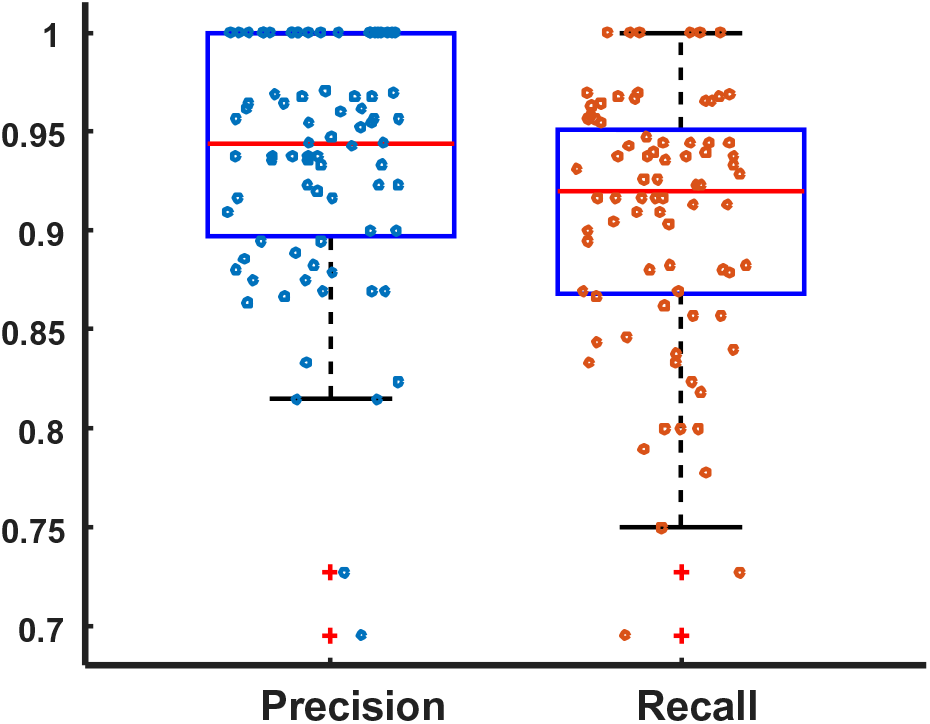
Performance of mitochondria detection by R-CNN. Evaluation of the mitochondrion test set in terms of precision and recall. Each circle indicates one EM image from the test set.

**Extended Data Figure 6.**
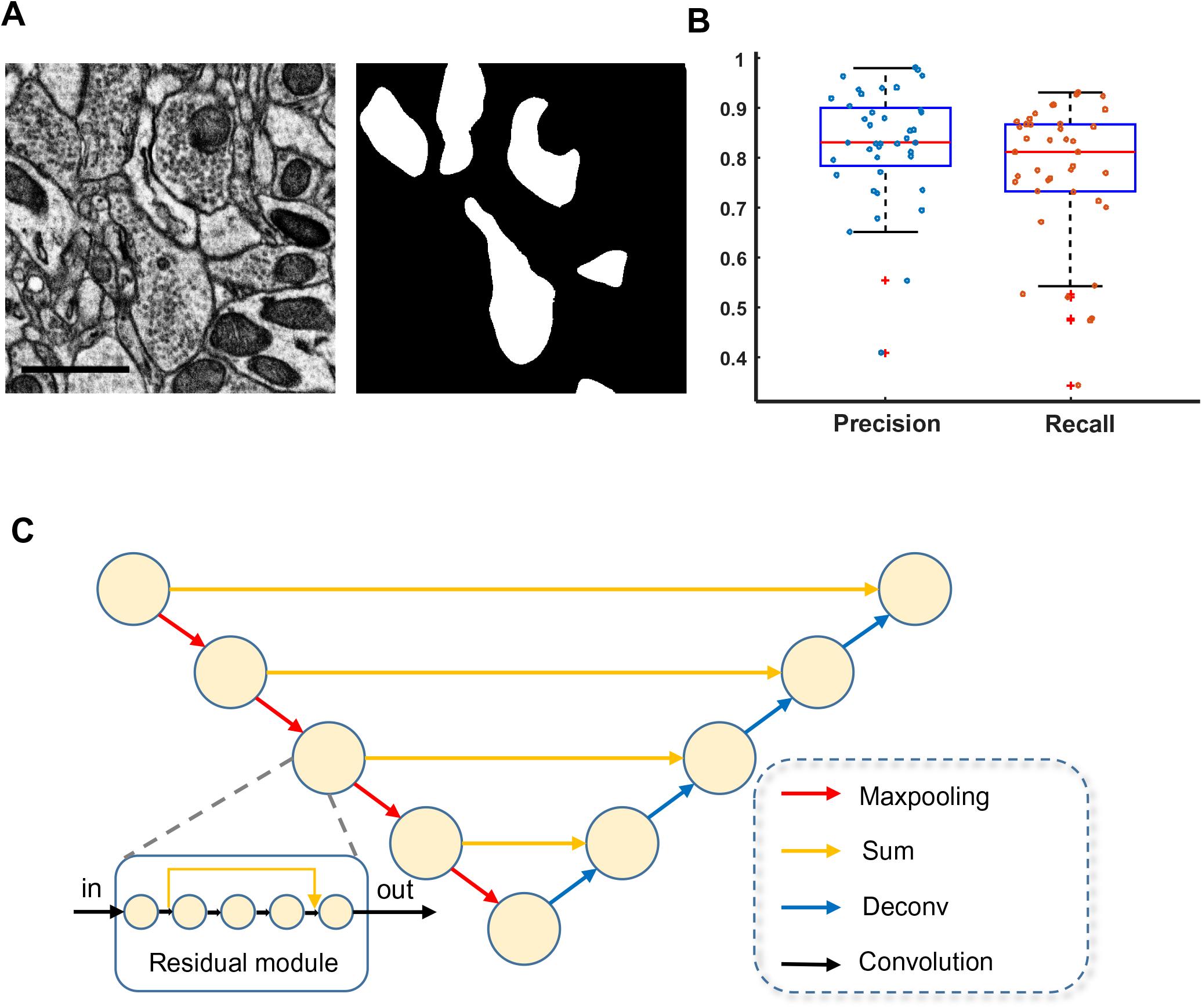
Identification of vesicle clouds. (A) Examples of EM image and binary vesicle cloud mask predicted by R-CNN. Scale bar: 1 μm. (B) Evaluation on the vesicle cloud test set in terms of precision and recall. Each circle indicates one EM image from the test set. (C) Network architecture of FusionNet, which takes a 1,024 × 1,024 image patch as input and outputs the binary vesicle cloud masks of input image. Each golden circle represents a residual module. The yellow arrows indicate the sum operation to fuse features from encode path and decode path. Red arrows and blue arrows indicate the convolution with maxpooling and deconvolution operations, respectively.

**Extended Data Figure 7.**
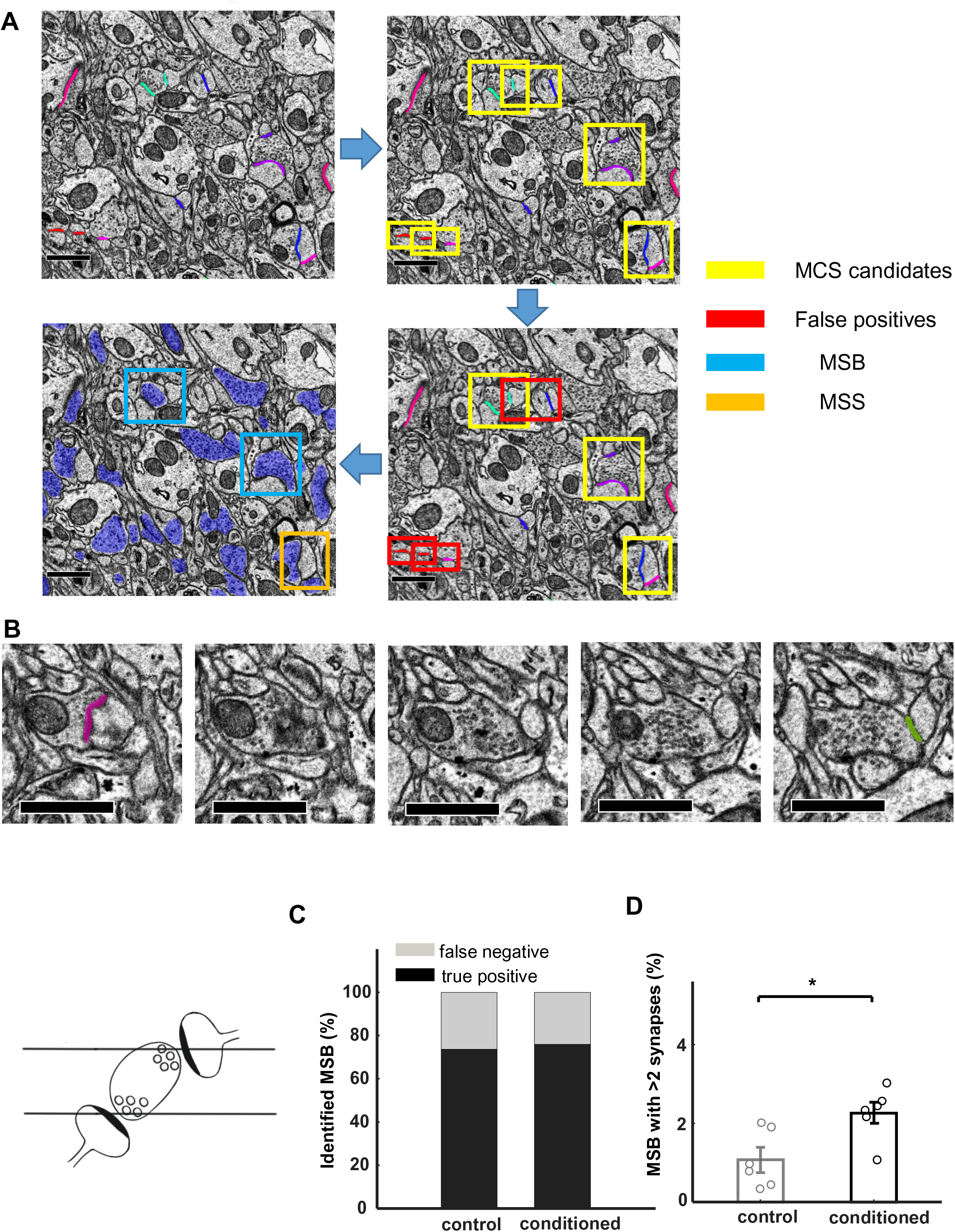
Semi-automatic detection, verification and classification of MCSs. (A) MCSs detection and verification. MCS candidates were first automatically marked based on a close-vicinity criterion, and then manually verified and classified into MSBs and MSSs based on the vesicle cloud features. Colored line: PSD; yellow box: MCS candidates; red box: false detection; blue box: verified MSB; orange box: verified MSS; blue patch: boutons. Scale bar: 1 μm. (B) A continuous series of EM images showing an MSB that was omitted by the automatic close-vicinity detection algorithm because the two spines never co-appeared on a single slice. Bottom left: a cartoon illustrating the false negative scenario. Scale bar: 1 μm. (C) False negative rate of MSBs estimated by manual identification of MSBs in saturated reconstruction blocks (control: 26.3%, tissue block 11 ×12 ×5 μm^3^, conditioned: 24%, 17 × 14 × 2.5 μm^3^). (D) The percentage of MSB synapses with more than 2 postsynaptic sites is higher in the conditioned mice (2.26 ± 0.3 %) than in controls (1.07 ± 0.3%). p < 0.05, two-sided t-test.

**Extended Data Figure 8.**
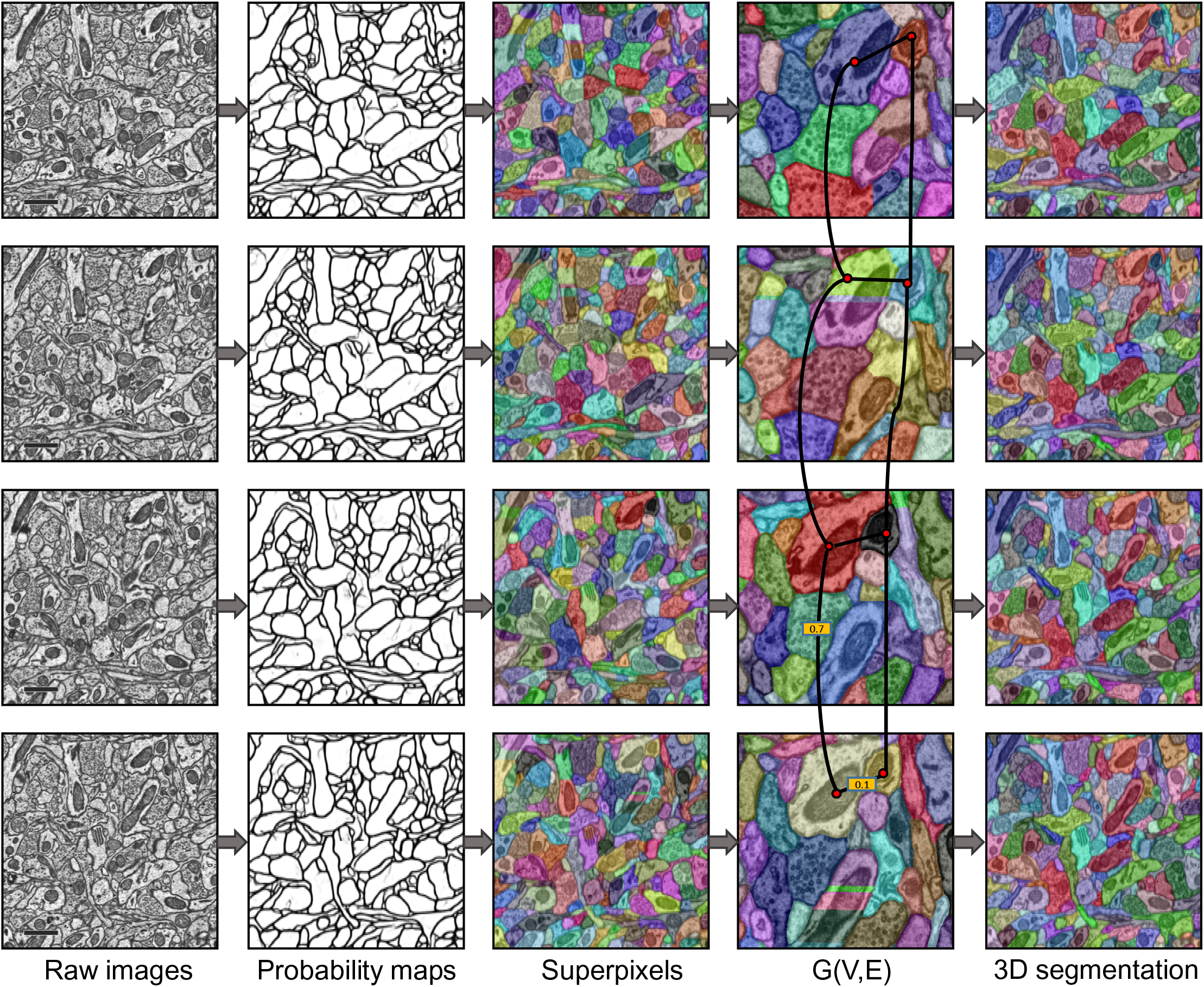
Workflow of volume segmentation. Trained network took raw images as input and membrane probability maps as output, then distance transform was applied to create superpixels and a graph G was constructed to build the relationships between segments. 3D segmentation results were obtained by solving the graph partitioning problem. Scale bar: 1 μm.

**Extended Data Figure 9.**
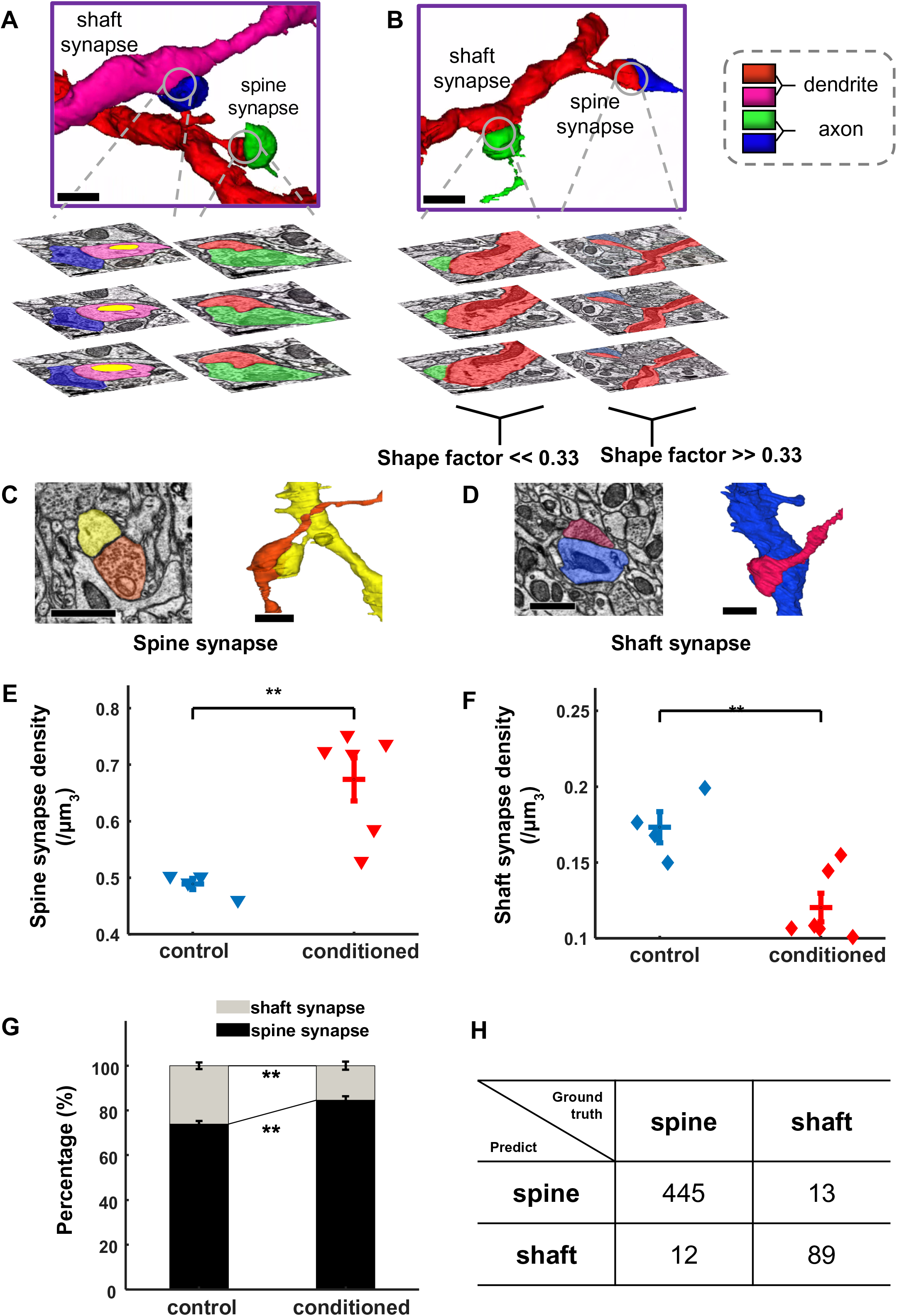
Classification of spine and shaft synapses. (A) Spine and shaft synapse classification based on the lack (right) or existence (left) of mitochondria (yellow) in postsynaptic structures. Scale bar: 1 μm. (B) Spine and shaft synapse classification based on spiny (right) or flat (left) shapes of postsynaptic structures. Scale bar: 1 μm. (C-D) Examples of a 3D reconstructed spine and a shaft synapse. Scale bar: 1 μm. (E) Spine synapse density is higher in the conditioned mice (0.674 ± 0.0379/µm^3^) than in controls (0.489 ± 0.0099/µm^3^). p<0.01, two-sided t-test. (F) Shaft synapse density is lower in the conditioned mice (0.12 ± 0.0094/µm^3^) than in controls (0.173 ± 0.0102/µm^3^). p<0.01, two-sided t-test. (G) Shaft and spine synapse proportion in the control mice (spine: 73.84 ± 1.49%, shaft: 26.16 ± 1.49%) and fear conditioned mice (spine: 84.54 ± 1.81%, shaft: 15.47 ± 1.81%) . p<0.01, two-sided t-test. (H) Confusion matrix of classification results on the test dataset.

**Extended Data Figure 10.**
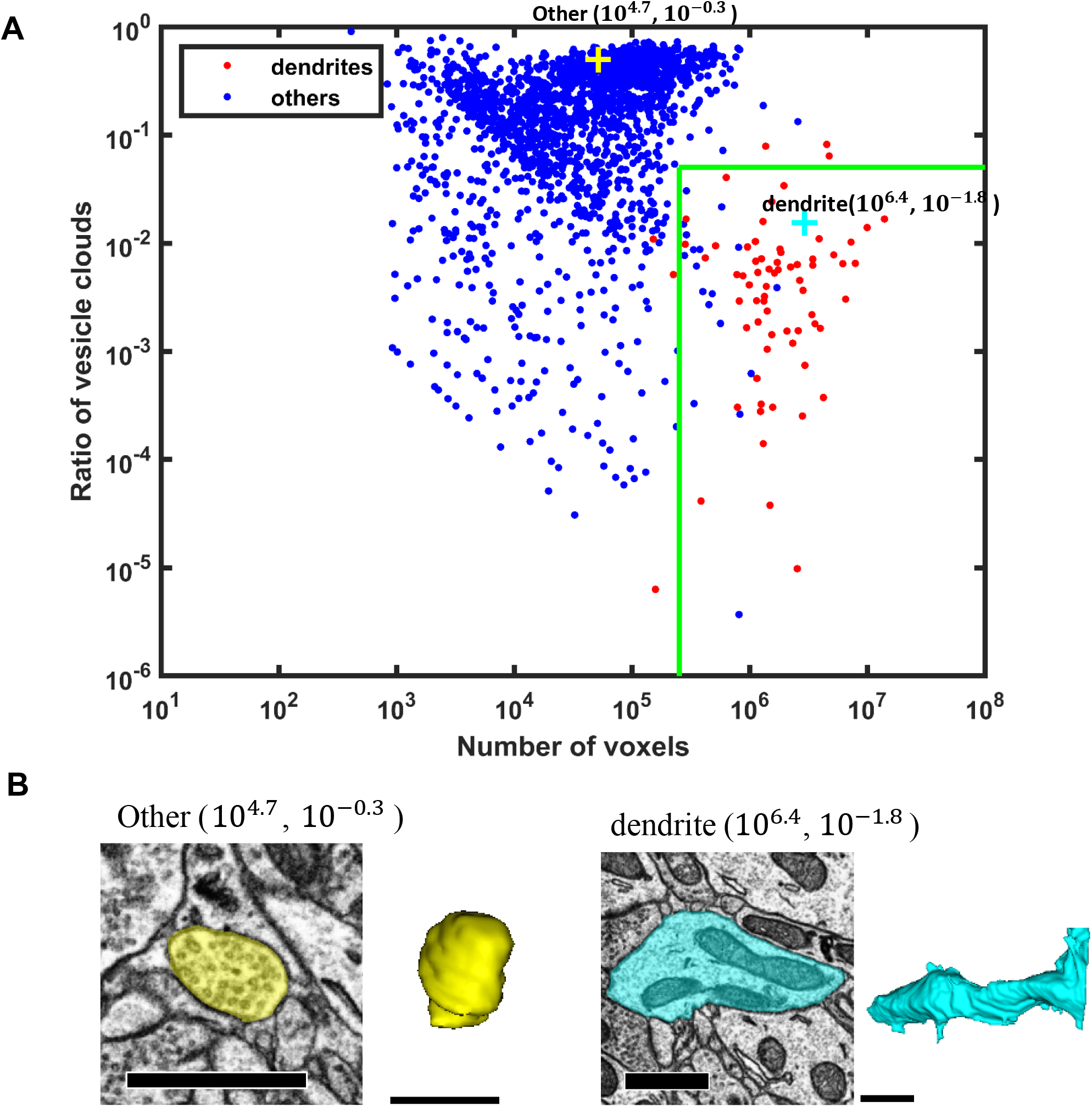
Identification of dendrites from saturated reconstruction. (A) Each point represents a dendrite or non-dendrite (other) segment. (B) Examples of identified axonal bouton (yellow) and dendritic branch (blue). Scale bar: 1 μm.

## Online Methods

### Animals

C57BL/6 mice were purchased from SLAC Laboratory Animals (Shanghai, China). YFP-H line mice were obtained from the Jackson Laboratory (Bar Harbor, ME, USA). Mice were bred and housed in the animal facility of Shanghai Protein Center under a 12 h light-dark cycle (7 am-7 pm light). Eight- to twelve-week-old male and female mice were used for the experiments. All procedures were approved by the Animal Committee of the Shanghai Tech University.

### Behavior

Fear conditioning and behavioral test for the freezing response took place in different environments. Mice were handled prior to conditioning. A commercial fear conditioning apparatus (MED Associates Inc., St. Albans, VT, USA) was used for the fear conditioning and behavioral test. Before conditioning and testing, the apparatus was wiped clean with 70% ethanol. The conditioned stimulus (CS) is a series of 14 kHz beeps (interleaved 0.5 s on, 0.5 s off) at 80 dB, lasting 10s in total; the unconditioned stimulus (US) is a 0.6 mA foot shock lasting 2 s. The sound co-terminates with the foot shock. For conditioned animals, CS-US pairing was presented 5 times with random intervals ranging from 60 to 90 s. For naïve animals, CS was presented 5 times with random intervals, without presentation of US. The behavioral responses to CS were tested 1 day after conditioning for both groups, using CS lasting 60 s. All conditioned animals showed high levels of freezing when CS was played, and naïve animals did not show a freezing response (Extended Data Figure 2).

### Electron microscopy sample preparation

The mouse was deeply anaesthetized with sodium pentobarbital (50 mg/kg, i.p.) and transcardially perfused with freshly prepared 4% paraformaldehyde and 0.5% glutaraldehyde (EM grade) in phosphate-buffered saline (PBS) 3 days after behavioral training. The brain was post-fixed in 2% paraformaldehyde and 2% glutaraldehyde overnight in a cold room. Blocks of the auditory cortex were dissected and embedded in resin for serial sectioning. Serial sections of the samples were continuously sectioned at 50 nm with an ATUMtome automated tape-collecting ultramicrotome (RMC, Boeckeler instruments Inc., Tucson, AZ, USA) and then collected onto a Kapton polyimide tape (8 mm wide and 100 μm thick). The tape with brain sections were then segmented and attached to 4-inch silicon wafers *via* double coated carbon conductive tape (TED Pella Inc, Redding, CA, USA). Lastly, the wafers were coated with 6 nm of carbon though an EM ACE60 high vacuum film deposition instrument (Leica Microsystems GmbH, Wetzlar, Germany) to prevent charging during scanning EM imaging (Figure 1).

### Electron Microscopy

The tape-collected ultra-thin sections were imaged on a Carl Zeiss Supra 55 scanning electron microscope (Carl Zeiss AG,. Oberkochen, Germany) using a secondary electron detection (9 kV accelerating potential, working distance of approximately 6.0 mm), with resolution of 2-4 nm/pixel and pixel dwelling time of 1.5 μs.

### Image Alignment

To correct for distortions of serial sections from automated tape-collecting ultramicrotome scanning electron microscopy (ATUM-SEM), we used a non-linear registration algorithm^38^ to create 3D image stacks that retain the original morphology as much as possible.

### Network Architecture of FPN

The FPN consists of a bottom-up pathway and a top-down pathway with lateral connections. On each level of the pyramid, the top-down pathway features will be fused (element-wise addition) with the bottom-up pathway features in the corresponding level. The scale-space induced from the feature pyramid fits well with the object-space. Thus, the RPN and R-CNN predict the class and regress the bounding box of objects at different scale using feature maps at different levels of the pyramid. The same applies for the mask branch.

### Block-wise inference strategy for large-scale data

Due to the constraints of the GPU, the inference was designed to proceed in a block-wise way for large-scale data (Extended Data Figure 3B). The original images were first cropped into small patches (2,048 × 2,048) with overlaps (100 × 100), which were then fed into the trained network to generate detection boxes and segmentation masks. To obtain the connection relationship at the 3D level, we used a strategy which first fused the results on 2D and then connected the adjacent 2D layers to produce the 3D results.

### 2D segmentation of neuronal processes

FusionNet^23^ was trained to predict the neuronal membrane. The membrane probability maps obtained from the network were binarized with a threshold of 0.5, and morphologically dilated with a disk radius of 2 in order to dismiss the small cracks in membranes and avoid merge errors. A watershed algorithm was then used to obtain connected neuronal components.

### Automated volume segmentation

The thickness of ssEM sections is a key factor for automatic reconstruction. High anisotropy brings more problems and challenges in learning the affinity between voxels along the z-direction. The state-of-the-art approach, which learns an affinity graph by 3D CNN, did not perform well on our data set. Therefore, we used the Multicut pipeline^33^ to obtain the volume segmentation. And we imported the results to the proofreading tool, and assigned the proofreading task to 6 experienced experimenters. The proofreading took about 4 weeks.

### Experimental setup

All networks were implemented in Keras with Tensorflow^39^ as backend. A stochastic gradient descent algorithm^40^ with learning rate of 0.001 was used to optimize the networks. To avoid overfitting, we used online data augmentation, including random rotation, random flipping and adding random noise for all training data sets. All training and inference procedures were performed on servers equipped with NVIDIA Tesla K40 GPUs.

### Spine and Shaft Synapse Classification

Excitatory and inhibitory synapses are classified according to some established criteria^41^. Due to the low axial resolution, indistinct synaptic vesicles and symmetry/asymmetry of PSDs can’t be used to identify the classes of synapses. Although it is generally believed that excitatory synapses are mostly located on spines, some studies have indicated that excitatory synapses can also form on dendritic shafts^42^, which cannot be quantified *in vivo* by counting spines using microscopy. Accordingly, based on the presence of postsynaptic mitochondria (Extended Data Figure 9A) and the shape of postsynaptic structures (Extended Data Figure 9B), we established some rules to classify the spine and shaft synapses (Extended Data Figure 9C and 9D).

### Error analysis of MCS detection algorithm

The multiple boutons or spines in a single MCS had to appear on the same section for them to be successfully identified by our semi-automatic MCS detection algorithm (Extended Data Figure 7B). Indeed, a comparison against manually quantified MSB/MSS suggested that the our semi-automatic MCS detection algorithm had a false negative detection rate of 25% for two densely reconstructed image stacks (Extended Data Figure 7C). The data in Figure 5D and Figure 5E were adjusted accordingly.

### Virus injection for *in vivo* imaging

The AAV-hSyn-EGFP and AAV-hSyn-tdTomato vectors were produced by Taitool Bioscience, Co., Ltd. (Shanghai, China). Virus injection was performed using a previously described protocol^3^. Briefly, mice were anaesthetized with sodium pentobarbital (7 mg/kg) and placed in a stereotaxic frame (RWD Life Sciences Co., Ltd., Shenzhen, China). For axonal labeling and imaging in the auditory cortex, 0.1-0.2 μl GFP or tdTomato viruses (∼10^13^ virus particles per ml) were injected, using a glass micropipette with a Nanoject III micro-injector (Drummond Scientific Company, Broomall, PA, USA), into three different regions that project axons to the auditory cortex: lateral amygdala (LA, 1.0 mm from Bregma, 3.25 mm lateral from the midline, 3.55 mm vertical from the cortical surface), anterior cingulate cortex (ACC, −1.00 mm from Bregma, 0.5 mm lateral from the midline, and 1.5 mm vertical from the cortical surface), medial geniculate nucleus: (MGN, 3.2 mm from bregma, 2.0 mm lateral from the midline, and 2.8 mm vertical from the cortical surface).

### Two-photon microscopy and data analysis

We performed cranial window implantation and two-photon imaging in mouse auditory cortex using previously described protocols^3^. Mice were imaged 1 day before and 3 days after fear conditioning. All images were analyzed using ImageJ (NIH, Bethesda, MD, USA). Dynamic turnover assays of boutons and spines were based on comparison of the images collected at two different time points (3 days after *vs*. 1 day before conditioning). Percentages were normalized to the initial image taken at 1 day before conditioning. Turnover of boutons was calculated by averaging the turnover rates of LA, ACC and MGN axonal boutons in the auditory cortex. For the dual-color labeled images, potential bouton-spine pairs were visually identified when a presynaptic bouton and a postsynaptic spine overlaid in the image stacks^3^.

### Mathematical modeling to assess bouton and spine turnover patterns: replacement/addition ratio

#### (1) Estimating bouton replacement and addition ratio by MSS percentages and bouton turnover rate

According to the ssEM data (Figure 4G), there were 1.8% MSS synapses in control animals and 1.4% in conditioned animals. Based on in vivo microscopic analysis (Figure 5B), the bouton elimination rate was approximately 15% and the formation rate was 15%. We built a model starting with 120 synapses consisting of 118 1-to-1 synapses (98.3%) and 2 MSS synapses (1 MSS; 1.7%, approximated to 1.8%), and ending with 120 synapses, also 118 1-to-1 synapses and 2 MSS synapse (1.7%, approximated to 1.4%). Note that in this model, the spine entities remained the same, and the number of boutons eliminated and the number newly formed were both 18 (120 × 15%) since the number of spines and MSSs were both constant.

To count all possible bouton turnover patterns, we considered bouton elimination before formation. There were a total of 3 types of bouton elimination, with the number of each type denoted as *a*_*k*_(*k* = 1,2,3) and the number of corresponding formation patterns satisfying the end situation denoted as *b*_*k*_(*k* = 1,2,3).

Thus, the total number of synaptic turnover patterns, denoted by *N*, can be calculated by:

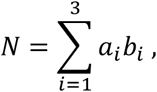

Then, we calculate the formation ratio of (A.), (B.), (D.), (E.), (F.) *vs*. (C.), indicating the percentage of bouton replacement and addition in new synapses, respectively.

#### (2) Estimating spine replacement and addition ratio by MSB percentages and spine turnover rate

According to the ssEM data (Figure 4F), 5.0 and 6.8% of all synapses were MSB synapses in control and conditioned animals, respectively. Based on in vivo imaging data (Figure 5B), spine turnover rate was approximately 10% for elimination and 10% for formation. The starting 120 synapses consisted of 114 1-to-1 synapses (95%) and 6 MSB synapses (3 MSBs; 5.0%), and the end situation consisted of 121 synapses including 12 new synapses (113 1-to-1 synapses and 8 MSB synapses, i.e., 4 MSBs, 6.6%, approximated to 6.8%). It should be noted that in this model, the bouton entities remain the same, and the difference between the number of synapses eliminated (11) and the number newly formed (120 × 10% = 12) is due to the constant number of boutons and variable number of MSBs.

To count all possible spine turnover patterns, we considered spine elimination before formation. There were a total of 10 types of spine elimination, with the number of each type denoted as *a*_*k*_′(*k* = 1,2, …, 10) and the number of corresponding formation patterns satisfying the end situation denoted as *b*_*k*_′(*k* = 1,2, …, 10).

Therefore, the total number of synaptic turnover patterns, denoted by *N*′, can be calculated by:

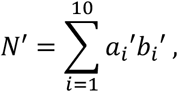

Then, we calculate the formation ratio of (A.), (B.), (D.), (E.), (F.) *vs.* (C.), indicating the percentage of spine replacement and addition in new synapses, respectively.

### Calculating the number of dendrites intersecting within 1-μm radius of a bouton

Based on the dense reconstruction results, we formulated some criteria to identify each neurite as dendritic or not. We extracted a 2D vector (number of voxels and ratio of vesicle cloud) as features to classify the segments. If the number of voxels in a neurite was greater than 250,000 and the ratio of vesicle cloud in it was less than 5%, this neurite was classified as dendritic (Extended Data Figure 10). For each bouton, we fit a minimum circumscribed circle and obtained the radius (R) and center (x, y, z) approximately. Then, we extended a sphere centered on (x, y, z) with radius of R+1 μm, and the number of dendrites that pass through the sphere was calculated.

### Static Synaptic Connectivity Model

For simplicity, our static model consists of 100 synaptic connections (Figure 6E). Based on our experimental data (Figure 6D), each bouton can make contact with its 9 potential dendrites. There are two notable constraints: 1) at most 2 synapses can be formed per bouton; 2) 6% of the synapses are MSB synapses (Figure 5G) for conditions 2 and 3.

For condition 1, the constraint of the number of synapses and only one synapse formed per bouton makes the existence of exactly 100 boutons. Each independent bouton has 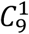 possible connections, so there is a total of 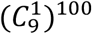 patterns for condition 1. For condition 2, since 6% of synapses are MSB synapses, there are a total of 6 MSB synapses from 3 MSBs. Thus, this condition consists of 97 boutons including 94 1-to-1 boutons and 3 MSBs. Accordingly, the number of patterns for condition 2 can be given by 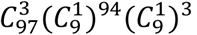. For condition 3, each of the 3 boutons selected as MSBs has 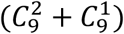 optional connection patterns that meet the condition. Rethinking the consideration of condition 2, the total number of patterns that meet condition 3 can be calculated as 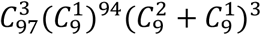.

### Plastic Synaptic Connectivity Model

We built a neural network model that incorporated plasticity by adding 10% more contacts to the boutons as a form of learning-induced synaptic formation. Plasticity is represented by a 10% increase in synaptic connections by adding 10 connections to existing boutons in a network consisting of 100 boutons and 100 1-to-1 synapses. There are two notable constraints: 1, at most one synapse is added to each bouton; 2, formation of the new synapses is random.

Before the synaptic formation, the number of possible connections for both conditions A and B was 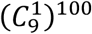, which is the same as condition 1 of the static model. For condition B, new connections can only be formed on the same dendrite. Therefore, only the factor of the selection of the 10 boutons from the 100 boutons to form new synapses can lead to an increase in the number of patterns triggered by the synaptic formation. Furthermore, the number of patterns after formation can be written as 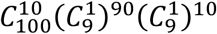. For condition A, like condition 3 in the static model, each of the 10 boutons selected as MSBs has 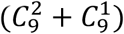 optional connection patterns that meet the criterion. Rethinking the consideration of condition B, the total number of patterns that meet condition A can be calculated as 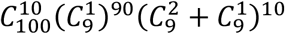.

## Footnotes

^1^ https://www.micro-visions.org/data/Synapse-ATUM/

