## Supplemental Methods for "Fear memory-associated synaptic and mitochondrial changes revealed by deep learning-based processing of electron microscopy data"

#### Data availability

The ultrastructural information of all reconstructed synapses was uploaded to a public website (https://www.micro-visions.org/data/Synapse-ATUM/database/) for future data mining.

#### Image Alignment

To correct for distortions of serial sections from automated tape-collecting ultramicrotome scanning electron microscopy (ATUM-SEM), we used a non-linear registration algorithm[1] to create 3D image stacks that retain the original morphology as much as possible. We assumed that the tissue deformations on different sections were independent. In order to extract reliable correspondences between adjacent sections, we used the dense correspondence matching SIFT-flow algorithm[2]. Then, the corresponding points on different sections were simultaneously adjusted based on an energy function to retain the same x-y coordinates. In addition, the displacements of these corresponding points were constrained to be smooth and small, thereby restricting the non-linear deformation of the original images. Finally, with the displacement vector of the extracted corresponding points, the positions of the points in the original sections, as well as in the aligned images, were obtained. The Moving-Least-Square (MLS) method[3] was used to warp each section image. The deformation result produced by the MLS method was globally smooth, and the biological tissue could retain its shape as a result of the rigid local transformation.

#### Network Architecture of FPN

The FPN consists of a bottom-up pathway and a top-down pathway with lateral connections. The bottom-up pathway leverages the hierarchy of the natural pyramid features of the ResNet50, and the top-down pathway recovers the original resolution by using the upsample layer step by step. On each level of the pyramid, the top-down pathway features will be fused (element-wise addition) with the bottom-up pathway features in the corresponding level. By constructing the feature pyramid, the lower-resolution feature maps contain more semantic information, while the higher-resolution feature maps contain more detailed information. The scale-space induced from the feature pyramid fits well with the object-space. Thus, the RPN and R-CNN predict the class and regress the bounding box of objects at different scale using feature maps at different levels of the pyramid. The same applies for the mask branch. Specifically, the feature pyramid has 5 levels and the corresponding object scales are 32, 64, 128, 256, 512, respectively.

#### Block-wise inference strategy for large-scale data

Due to the constraints of the GPU, the inference was designed to proceed in a block-wise way for large-scale data (Extended Data Figure 3B). The original images were first cropped into small patches ($2{,}048 \times 2{,}048$) with overlaps ($100 \times 100$), which were then fed into the trained network to generate detection boxes and segmentation masks. To obtain the connection relationship at the 3D level, we used a strategy which first fused the results on 2D and then connected the adjacent 2D layers to produce the 3D results.

To facilitate the description of our algorithm, we defined $N_i$ as the number of detection boxes in the i-th layer, and $D_{ij}$ represented the j-th synaptic detection box in the i-th layer:

$$D_{ij} = (x_{ij}^1, y_{ij}^1, x_{ij}^2, y_{ij}^2)$$

where $x_{ij}^1, y_{ij}^1, x_{ij}^2, y_{ij}^2$ represented the left-upper coordinate, left-upper coordinate, right-lower coordinate and right-lower coordinate, respectively; $D_i = \{D_{ij},\ j = 1,2, \dots, N_i\}$ denoted the set of all detection boxes on the i-th layer. $C_{ij}$ represented the corresponding segmentation result of the j-th synapse in the i-th layer. $C_i = \{C_{ij},\ j = 1,2, \dots, N_i\ \}$ represented the set of all binary segmentation results in the i-th layer.

**（1）Single-layer fusion algorithm for detection and segmentation results**

After obtaining the synaptic detection and the segmentation results of small patches by deep neural network, we stitched them to recover the original image size. At the overlapping region, there could be multiple different detection results for the same synapse. In this case, the synapse was repeatedly detected. Therefore, we designed an iterative 2D fusion algorithm to fuse the detection bounding boxes and corresponding segmentation masks in the overlapping areas. The main procedures are as follows:

Step 1: Construct the Intersection-over-Union (IoU) matrix $S_i$ between all candidate detection boxes $D_{ij}$ in the i-th layer. The $S_i$ matrix can be formulated as:

$$S_i = \begin{pmatrix} s_{11}^i & s_{12}^i & \cdots & s_{1N_i}^i \\ s_{21}^i & s_{22}^i & \cdots & s_{2N_i}^i \\ \vdots & \vdots & \ddots & \vdots \\ s_{N_i1}^i & s_{N_i2}^i & \cdots & s_{N_iN_i}^i \end{pmatrix}$$

where $s_{jk}^i$ represents the IoU of the j-th detection box and the k-th detection box, and the calculation formula can be expressed as follows:

$$s_{jk}^i = \frac{A(\cap \{D_{ij}, D_{ik}\})}{A(\cup \{D_{ij}, D_{ik}\})}, j = 1,2, \dots, N_i$$

where $A$ is a function of calculating area. $\cap \{D_{ij}, D_{ik}\}$ and $\cup \{D_{ij}, D_{ik}\}$ represent the intersection and union of $D_{ij}$ and $D_{ik}$, respectively. Obviously, $s_{jk}^i \epsilon [0,1]$. It is not difficult to find that $S_i$ is a sparsely symmetric matrix with a main diagonal of 1.

Step 2: Find the coordinates of elements in $S_i$, which are greater than the threshold $\theta_1$. Let $Ind$ be a set, then:

$$Ind = \text{II}_{\{S_i > \theta_1\}}$$

where II denotes the indicator function. This operation acts on each element of $S_i$ and returns the row and column number $jk$ satisfying the condition.

Step 3: For $\forall jk \in Ind$, merge $D_{ij}$ and $D_{ik}$ into a new detection box $D'_{ijk}$. The coordinates of the new detection box are illustrated as follows:

$$D'_{ijk} = (\min(x^1_{ij}, x^1_{ik}), \min(y^1_{ij}, y^1_{ik}), \max(x^2_{ij}, x^2_{ik}), \max(y^2_{ij}, y^2_{ik}))$$

Step 4: Update $N_i$ and $D_i$, and repeat above steps until no box need to be merged.

**（2）Multilayer 3D connection algorithm**

In order to obtain 3D synapses from the serial 2D segmentation results as well as screen out false positives, we develop and implement a 3D connection algorithm in the fine registered stacks according to the continuity of ssEM images and the spatial structural information of synapses. This procedure not only recovers synaptic morphology at the 3D level, but also assigns a unique label to each 3D synapse. The main steps are as follows:

Step 1: Construct the similarity matrix $S^{it}$ between the synapses in the i-th layer and synapses in the t-th layer. $S^{it}$ can be formulated as follows:

$$S^{it} = \begin{pmatrix} s^{it}_{11} & s^{it}_{12} & \cdots & s^{it}_{1N_i} \\ s^{it}_{21} & s^{it}_{22} & \cdots & s^{it}_{2N_i} \\ \vdots & \vdots & \ddots & \vdots \\ s^{it}_{N_i1} & s^{it}_{N_i2} & \cdots & s^{it}_{N_iN_i} \end{pmatrix}$$

where t = i+1. It should be noted that $s^{it}_{pq}$ represents the ratio of the overlapping area between $D_{ip}$ and $D_{tq}$ to the minimum area of these two, and the calculation formula is:

$$s^{it}_{pq} = \frac{A(\cap\{D_{ip}, D_{tq}\})}{\min(A(D_{ip}), A(D_{tq}))}$$

Step 2: Find the coordinates of elements of $S^{it}$, which are greater than the threshold $\theta_2$, *i.e.*

$$Ind1 = \mathrm{II}_{\{S^{it} > \theta_2\}}$$

where II denotes the indicator function. This operation acts on each element of $S^{it}$ and returns the row and column number $pq$ satisfying the condition.

Step 3: For $\forall pq \in Ind1$, assign the same and unique label value to $C_{tq}$ and $C_{ip}$. Then, the binary segmentation is converted into a label image.

Step 4: Repeat above steps for all layers and obtain a labeled stack. Look through all the labels to check the number of layers $L^t$ for each label $t$. If $L^t$ is less than the predefined threshold $\theta_3$, then delete the corresponding segmentation $C_{it}$ from the original results, which can be expressed as follows:

$$C_i = C_i \backslash C_{it}$$

After implementing the block-wise algorithm, we obtain the final result of each large image, where the same label value indicates the same synapse at the 3D level.

#### 2D segmentation of neuronal processes

We extracted two volumes (2,048 × 2,048 × 50 voxels) from control and conditioned group as the training data for membrane detection and dense reconstruction. In the ground truth, voxels with the same value belong to the same neurite in 3D. Three experienced annotators labeled the dense labels with cross-validation.

FusionNet[4] was trained to predict the neuronal membrane. The training data was extracted from the volume segmentation data set. Pixels that were labeled as background or at the edge of any adjacent neurite objects were collected as boundaries. The membrane probability maps obtained from the network were binarized with a threshold of 0.5, and morphologically dilated with a disk radius of 2 in order to dismiss the small cracks in membranes and avoid merge errors. A watershed algorithm was then used to obtain connected neuronal components.

#### Automated volume segmentation

The thickness of ssEM sections is a key factor for automatic reconstruction. High anisotropy brings more problems and challenges in learning the affinity between voxels along the z-direction. The state-of-the-art approach, which learns an affinity graph by 3D CNN, did not perform well on our data set. Therefore, we used the Multicut pipeline[5] to analyze our data. We first applied a distance transform to generate superpixels in 2D slices. Subsequently, we constructed a 3D region adjacency graph to connect the superpixels in 2D slices as well as between sections. We then abstracted the graph as G(V, E), where the node set V denoted all the superpixels and the edge set E represented boundaries between adjacent superpixels. Then, a random forest classifier was trained to predict the scores (probability of whether an edge should be cut) of every edge in the graph to obtain a weighted un-directed graph (Extended Data Figure 8). We thus solved the graph partitioning problem with an approximate solution. Finally, we imported the segmentation results to the proofreading tool, and assigned the proofreading task to 6 experienced experimenters. The proofreading took about 4 weeks.

#### Spine and Shaft Synapse Classification

Excitatory and inhibitory synapses are classified according to some established criteria[6]. Due to the low axial resolution, indistinct synaptic vesicles and symmetry/asymmetry of PSDs can't be used to identify the classes of synapses. Although it is generally believed that excitatory synapses are mostly located on spines, some studies have indicated that excitatory synapses can also form on dendritic shafts[7], which cannot be quantified *in vivo* by counting spines using microscopy. Accordingly, based on the presence of postsynaptic mitochondria (Extended Data Figure 9A) and the shape of postsynaptic structures (Extended Data Figure 9B), we established some rules to classify the spine and shaft synapses (Extended Data Figure 9C and 9D).

Previous studies have shown that there are very few mitochondria in dendritic spines, thus the absence of mitochondria can be used as one informative feature for spine identification. If no mitochondria are present in the postsynaptic site on any layer, the synapse is classified as a "spine synapse". If the proportion of mitochondria present on all layers is greater than 50%, the synapse is classified as a "shaft synapse". Since the sectioning orientation may lead to false negatives (*e.g.*, Extended Data Figure 9B), we added a morphological criterion, *i.e.*, whether the shape of the postsynaptic element is spiny or flat, for classification of remaining conditions. The mean change rate of postsynaptic areas from adjacent sections is defined as the shape factor—another informative feature for spine identification. A higher shape factor value (greater than

0.33) indicates a higher probability of existence of spines; a lower shape factor value indicates the presence of shafts (Extended Data Figure 9B). The evaluation on the test data set consisting of 559 synapses showed that our method yielded an accuracy rate of 0.95 (Extended Data Figure 9H).

#### Details for Mathematical modeling to assess bouton and spine turnover patterns: replacement/addition ratio

##### (1) Estimating bouton replacement and addition ratio by MSS percentages and bouton turnover rate

According to the ssEM data (Figure 4G), there were 1.8% MSS synapses in control animals and 1.4% in conditioned animals. Based on *in vivo* microscopic analysis (Figure 5B), the bouton elimination rate was approximately 15% and the formation rate was 15%. We built a model starting with 120 synapses consisting of 118 1-to-1 synapses (98.3%) and 2 MSS synapses (1 MSS; 1.7%, approximated to 1.8%), and ending with 120 synapses, also 118 1-to-1 synapses and 2 MSS synapse (1.7%, approximated to 1.4%). Note that in this model, the spine entities remained the same, and the number of boutons eliminated and the number newly formed were both 18 (120 $\times$ 15%) since the number of spines and MSSs were both constant. Three major assumptions were made: 1) each bouton forms at most one synapse (MSB is not considered in this model); 2) each spine contains 1 or 2 synapses (for simplicity of modeling, we do not consider MSS containing 3 or more synapses); 3) the elimination/formation of synapses is represented by the elimination/formation of boutons.

To count all possible bouton turnover patterns, we considered bouton elimination before formation. There were a total of 3 types of bouton elimination, with the number of each type denoted as $a_k (k = 1,2,3)$ and the number of corresponding formation patterns satisfying the end situation denoted as $b_k (k = 1,2,3)$. The three types are as follows:

(1) All 18 bouton eliminations are from 1-to-1 synapses (118 in total), $a_1 = C_{118}^{18}$; All 18 bouton formations occur on above alone spines, $b_1 = 1$;
(2) There are 17 eliminations from 1-to-1 synapses and 1 from MSS synapses (2 in total), $a_2 = C_{118}^{17} C_2^1$; 17 bouton formations on above alone spines and then another one on one of the 119 candidate 1-to-1 synapses (118-17+1+17) to form MSS, $b_2 = C_{119}^1$;
(3) There are 16 eliminations from 1-to-1 synapses and 2 eliminations from 1 MSS, $a_3 = C_{118}^{16} C_2^2$; 17 bouton formations on above alone spines and then another one on one of the 119 candidate 1-to-1 synapses (118-16+17) to form MSS, $b_3 = C_{119}^1$.

Thus, the total number of synaptic turnover patterns, denoted by $N$, can be calculated by:

$$N = \sum_{i=1}^{3} a_i b_i ,$$

Then, we calculate the formation ratio of (A.), (B.), (D.), (E.), (F.) *vs*. (C.), indicating the percentage of bouton replacement and addition in new synapses, respectively.

Assuming that the possibility of each synaptic turnover pattern is equal, the mathematical expectation for the number of synapses corresponding to Situation (C.) in 18 new synapses, $n_{add}$, and the corresponding number of new synapses where bouton replacement occurs, $n_{replace}$, can be derived using the following equations:

$$n_{add} = \frac{\sum_{j=0}^{18} jN_j}{N'},$$

$$n_{replace} + n_{add} = 18,$$

where, $N_j (j = 0,1,2,\ldots,18)$ is the number of synaptic turnover patterns in which only $j$ out of 18 new synapses belong to Situation (C.). As the number of MSS after learning is only one, there can be no more than 1 addition. So $N_2 = N_3 = \cdots = N_{17} = N_{18} = 0$. $N_j (j = 0,1)$ can be obtained as follows:

$$N_0 = N - \sum_{j=1}^{18} N_j,$$

$$N_1 = a_2 C_{101}^1 + a_3 C_{102}^1.$$

In order to gain a better understanding of the above formula, both items in $N_1$ are elaborated as follows. For elimination type (2), after adding one synapse to each of the 17 alone spines, the last formed synapse to be determined is randomly added to one of 101 1-to-1 synapses that have previously remained unchanged (118-17), so the number of patterns is calculated as $a_2 C_{101}^1$. For elimination type (3), the last formed synapse to be determined is randomly added to one of 102 1-to-1 synapses that have previously remained unchanged (118-16), and thus the number of patterns is obtained as $a_3 C_{102}^1$.

Using the above equations, we determined that of the 18 new boutons, on average, 17.17 were accompanied by old bouton elimination (*i.e.*, replacement) and the other 0.83 were added to existing 1-to-1 synapses to form MSSs, accounting for 95.4 and 4.6%, respectively.

In addition, we extended the model programmatically to any size regarding the number of synapses. The input parameters of the program included only the ratio of multiple-contact synapses before and after learning, turnover rate, and the number of synapses. To test our model with different parameters: the last parameter is scaled up to 12,000 and the other parameters are fixed. In general, the obtained results with different model sizes were similar, as shown in Figure 5G and Table 1.

**Table 1. the bouton replacement and addition ratio with different model sizes**

| Model size | 120 | 1,200 | 12,000 |
|---|---|---|---|
| $n_{replace}$ (%) | 95.4% | 96.3% | 96.1% |
| $n_{add}$ (%) | 4.6% | 3.7% | 3.9% |

##### (2) Estimating spine replacement and addition ratio by MSB percentages and spine turnover rate

According to the ssEM data (Figure 4F), 5.0 and 6.8% of all synapses were MSB synapses in control and conditioned animals, respectively. Based on *in vivo* imaging data (Figure 5B), spine turnover rate was approximately 10% for elimination and 10% for formation. The starting 120 synapses consisted of 114 1-to-1 synapses (95%) and 6 MSB synapses (3 MSBs; 5.0%), and the end situation consisted of 121 synapses including 12 new synapses (113 1-to-1 synapses and 8 MSB synapses, *i.e.*, 4 MSBs, 6.6%, approximated to 6.8%). It should be noted that in this model, the bouton entities remain the same, and the difference between the number of synapses eliminated (11) and the number newly formed (120 × 10% = 12) is due to the constant number of boutons and variable number of MSBs. Three major assumptions were made: 1) each spine forms at most one synapse (MSS is not considered in this model); 2) each bouton contains 1 or 2 synapses; 3) the elimination/formation of synapses is represented by the elimination/formation of spines.

To count all possible spine turnover patterns, we considered spine elimination before formation. There were a total of 10 types of spine elimination, with the number of each type denoted as $a_k'(k = 1,2,\ldots,10)$ and the number of corresponding formation patterns satisfying the end situation denoted as $b_k'(k = 1,2,\ldots,10)$. The 10 types are as follows:

(1') All 11 spine eliminations are from 1-to-1 synapses (114 in total), $a_1' = C_{114}^{11}$; 11 out of 12 spine formations occur on above alone boutons and then another one on one of the 114 candidate 1-to-1 synapses (114-11+11) to form MSB, $b_1' = C_{114}^{1}$;

(2’) There are 10 eliminations from 1-to-1 synapses and 1 from MSB synapses (6 in total), $a_2{}' = C_{114}^{10}C_3^1C_2^1$; 10 spine formations on above alone boutons and then another two on 115 1-to-1 synapses (114-10+1+10), $b_2{}' = C_{115}^2$;

(3’) There are 9 eliminations from 1-to-1 synapses and 2 eliminations from one single MSB, $a_3{}' = C_{114}^9C_3^1C_2^2$; 10 spine formations on above alone boutons and then another two on 115 1-to-1 synapses (114-9+10), $b_3{}' = C_{115}^2$;

(4’) 9 eliminations from 1-to-1 synapses and 2 eliminations from two different MSBs, $a_4{}' = C_{114}^9C_3^2(C_2^1)^2$; 9 spine formations on above alone boutons and then another three on 116 1-to-1 synapses (114-9+2+9), $b_4{}' = C_{116}^3$;

(5’) There are 8 eliminations from 1-to-1 synapses and 3 eliminations from MSBs, of which 2 are on the same MSB, $a_5{}' = C_{114}^8C_3^1C_2^2C_2^1C_2^1$; 9 spine formations on above alone boutons and then another three on 116 1-to-1 synapses (114-8+1+9), $b_5{}' = C_{116}^3$;

(6’) There are 8 eliminations from 1-to-1 synapses and 3 eliminations from different MSBs, $a_6{}' = C_{114}^8C_3^3(C_2^1)^3$; 8 spine formations on above alone boutons and then another four on 117 1-to-1 synapses (114-8+3+8), $b_6{}' = C_{117}^4$;

(7’) There are 7 eliminations from 1-to-1 synapses and 4 eliminations from 2 MSBs, $a_7{}' = C_{114}^7C_3^2(C_2^2)^2$; 9 spine formations on above alone boutons and then another three on 116 1-to-1 synapses (114-7+9), $b_7{}' = C_{116}^3$;

(8’) There are 7 eliminations from 1-to-1 synapses and 4 eliminations from 3 MSBs, $a_8{}' = C_{114}^7C_3^1C_2^2C_2^2(C_2^1)^2$; 8 spine formations on above alone boutons and then another four on 117 1-to-1 synapses (114-7+2+8), $b_8{}' = C_{117}^4$;

(9’) There are 6 eliminations from 1-to-1 synapses and 5 eliminations from 3 MSBs,

$a_9{}' = C_{114}^6 C_3^2 (C_2^2)^2 C_2^1$; 8 spine formations on above alone boutons and then another four on 117 1-to-1 synapses (114-6+1+8), $b_9{}' = C_{117}^4$;

(10’) There are 5 eliminations from 1-to-1 synapses and 6 eliminations from 3 MSBs, $a_{10}{}' = C_{114}^5 C_3^3 (C_2^2)^3$; 8 spine formations on above alone boutons and then another four on 117 1-to-1 synapses (114-5+8), $b_{10}{}' = C_{117}^4$.

Therefore, the total number of synaptic turnover patterns, denoted by $N'$, can be calculated by:

$$N' = \sum_{i=1}^{10} a_i{}' b_i{}' \,,$$

Then, we calculate the formation ratio of (A.), (B.), (D.), (E.), (F.) *vs.* (C.), indicating the percentage of spine replacement and addition in new synapses, respectively. Assuming that the possibility of each synaptic turnover pattern is equal, the mathematical expectation for the number of synapses corresponding to Situation (C.) in 12 new synapses, $n_{add}{}'$, and the corresponding number of new synapses where spine replacement occurs, $n_{replace}{}'$, can be derived from the following equations:

$$n_{add}{}' = \frac{\sum_{j=0}^{12} j N_j{}'}{N'} \,,$$

$$n_{replace}{}' + n_{add}{}' = 12 \,,$$

where, $N_j{}'(j = 0,1,2, \ldots ,12)$ is the number of synaptic turnover patterns in which only $j$ out of 12 new synapses belong to Situation (C.). As the number of MSBs after learning is 4, there can be no more than 4 additions. So, $N_5' = N_6' = \cdots = N_{11}' = N_{12}' = 0$. $N_j'$ $(j = 0,1,\ldots,4)$ can be obtained as follows:

$$N_0' = N' - \sum_{j=1}^{12} N_j' ,$$

$$N_1' = a_1' C_{103}^1 + a_2' C_{104}^1 C_{11}^1 + a_3' C_{105}^1 C_{10}^1 + a_4' C_{105}^1 C_{11}^2 + a_5' C_{106}^1 C_{10}^2 + a_6' C_{106}^1 C_{11}^3$$

$$+ a_7' C_{107}^1 C_9^2 + a_8' C_{107}^1 C_{10}^3 + a_9' C_{108}^1 C_9^3 + a_{10}' C_{109}^1 C_8^3 ,$$

$$N_2' = a_2' C_{104}^2 + a_3' C_{105}^2 + a_4' C_{105}^2 C_{11}^1 + a_5' C_{106}^2 C_{10}^1 + a_6' C_{106}^2 C_{11}^2 + a_7' C_{107}^2 C_9^1$$

$$+ a_8' C_{107}^2 C_{10}^2 + a_9' C_{108}^2 C_9^2 + a_{10}' C_{109}^2 C_8^2 ,$$

$$N_3' = a_4' C_{105}^3 + a_5' C_{106}^3 + a_6' C_{106}^3 C_{11}^1 + a_7' C_{107}^3 + a_8' C_{107}^3 C_{10}^1 + a_9' C_{108}^3 C_9^1 + a_{10}' C_{109}^3 C_8^1 ,$$

$$N_4' = a_6' C_{106}^4 + a_8' C_{107}^4 + a_9' C_{108}^4 + a_{10}' C_{109}^4 .$$

The first three items in $N_1'$ are selected for detailed explanation. For elimination type (1'), after adding one synapse to each of the 11 alone boutons, the last formed synapse to be determined is randomly added to one of 103 1-to-1 synapses that have previously remained unchanged (114-11), and therefore the number of patterns is obtained as $a_1' C_{103}^1$. For elimination type (2'), the last two newly formed synapses to be determined are randomly added to one of 104 1-to-1 synapses that have previously remained unchanged (114-10) and one of other 11 "new" 1-to-1 synapses (10 1-to-1 synapses formed by 1 elimination of old 1-to-1 synapses and subsequent 1 addition; 1 1-to-1 synapse by eliminating 1 synapse of MSB), respectively. So, the number of patterns is calculated as $a_2' C_{104}^1 C_{11}^1$. For elimination type (3'), the last two new synapses to be determined are randomly added to one of 105 1-to-1 synapses that have previously remained unchanged (114-9) and one of other 10 "new" 1-to-1 synapses (9 formed by

1 elimination of old 1-to-1 synapses and subsequent 1 addition; 1 formed by 2 synaptic elimination of an old MSB and subsequent 1 addition), respectively. So, the number of patterns is obtained as $a_3{}'C_{105}^{1}C_{10}^{1}$.

Using the above equations, we determined that of the 12 new spines, on average, 8.78 were accompanied by old spine elimination (i.e., replacement) and the other 3.22 were added to existing 1-to-1 synapses to form MSBs, accounting for 73.2 and 26.8%, respectively.

In addition, we also developed the model programmatically for any model size about the number of synapses. In general, the obtained results with different model sizes are similar, as shown in Figure 5H and Table 2.

**Table 2. the spine replacement and addition ratio with different model sizes**

| Model size | 120 | 1,200 | 12,000 |
| --- | --- | --- | --- |
| $n_{replace}$ (%) | 73.2% | 72.6% | 72.8% |
| $n_{add}$ (%) | 26.8% | 27.4% | 27.2% |

### Details for Static Connectivity Model

For simplicity, our model consists of 100 synaptic connections (Figure 6E). Based on our experimental data (Figure 6D), each bouton can make contact with its 9 potential dendrites. There are two notable constraints: 1) at most 2 synapses can be formed per bouton; 2) 6% of the synapses are MSB synapses (Figure 5G) for conditions 2 and 3.

For condition 1, the constraint of the number of synapses and only one synapse formed per bouton makes the existence of exactly 100 boutons. Each independent bouton has $C_9^1$ possible connections, so there is a total of $(C_9^1)^{100}$ patterns for condition 1.

For condition 2, since 6% of synapses are MSB synapses, there are a total of 6 MSB synapses from 3 MSBs. Thus, this condition consists of 97 boutons including 94 1-to-1 boutons and 3 MSBs. Accordingly, the number of patterns for condition 2 can be given by $C_{97}^3(C_9^1)^{94}(C_9^1)^3$. In other words, the selection of 3 MSBs from 97 boutons and the independent connection of each bouton are all factors that contribute to the increase of the number of patterns.

For condition 3, each of the 3 boutons selected as MSBs has $(C_9^2 + C_9^1)$ optional connection patterns that meet the condition. Rethinking the consideration of condition 2, the total number of patterns that meet condition 3 can be calculated as $C_{97}^3(C_9^1)^{94}(C_9^2 + C_9^1)^3$.

In detail, we express the respective calculation equations of information entropy that satisfy conditions 1, 2 and 3, as follows:

$$H_{condition1} = -\sum_{i=1}^{(C_9^1)^{100}} \frac{1}{(C_9^1)^{100}} log_2 \frac{1}{(C_9^1)^{100}} = 317bits\,,$$

$$H_{condition2} = -\sum_{i=1}^{C_{97}^3(C_9^1)^{97}} \frac{1}{C_{97}^3(C_9^1)^{97}} log_2 \frac{1}{C_{97}^3(C_9^1)^{97}} = 325bits\,,$$

$$H_{condition3} = -\sum_{i=1}^{C_{97}^{3}\left(C_{9}^{1}\right)^{94}\left(C_{9}^{2}+C_{9}^{1}\right)^{3}} \frac{1}{C_{97}^{3}(C_{9}^{1})^{94}(C_{9}^{2}+C_{9}^{1})^{3}} log_2 \frac{1}{C_{97}^{3}(C_{9}^{1})^{94}(C_{9}^{2}+C_{9}^{1})^{3}} = 332\ bits\ ,$$

where, $C_n^m$ is the combinatorial number with respect to $n$ and $m$.

Thus, including an MSB that connect to the same dendrite increases the ISC by 2.5% in this model, and the connectivity of the MSB to multiple dendrites increases another 2.2%. The results hold when we scale up the model tenfold each time up to $10^6$ synapses (Figure 6G). The information entropy values of 3 conditions under 5 model scales are listed in Table 3. The benefits of adding MSBs remained when the model was scaled up to $10^6$ synapses (Figure 6G).

**Table 3. The information storage capacity (ISC) of a static neural network**

| Model scale | $10^2$ | $10^3$ | $10^4$ | $10^5$ | $10^6$ |
|---|---|---|---|---|---|
| Condition 1 | 317 bits | 3,170 bits | 31,699 bits | 316,993 bits | 3,169,925 bits |
| Condition 2 | 325 bits | 3,264 bits | 32,673 bits | 326,781 bits | 3,267,872 bits |
| Condition 3 | 332 bits | 3,334 bits | 33,370 bits | 333,747 bits | 3,337,529 bits |

### Details for Plastic Connectivity Model

We built a neural network model that incorporated plasticity by adding 10% more contacts to the boutons as a form of learning-induced synaptic formation. Plasticity is represented by a 10% increase in synaptic connections by adding 10 connections to existing boutons in a network consisting of 100 boutons and 100 1-to-1 synapses. There are two notable constraints: 1, at most one synapse is added to each bouton; 2, formation of the new synapses is random.

Before the synaptic formation, the number of possible connections for both conditions A and B was $(C_9^1)^{100}$, which is the same as condition 1 of the static model.

For condition B, new connections can only be formed on the same dendrite. Therefore, only the factor of the selection of the 10 boutons from the 100 boutons to form new synapses can lead to an increase in the number of patterns triggered by the synaptic formation. Furthermore, the number of patterns after formation can be written as $C_{100}^{10}(C_9^1)^{90}(C_9^1)^{10}$.

For condition A, like condition 3 in the static model, each of the 10 boutons selected as MSBs has $(C_9^2 + C_9^1)$ optional connection patterns that meet the criterion. Rethinking the consideration of condition B, the total number of patterns that meet condition A can be calculated as $C_{100}^{10}(C_9^1)^{90}(C_9^2 + C_9^1)^{10}$. We can obtain the increase of information entropy for conditions A and B, as follows:

$$\Delta H_{conditionA} = -\sum_{i=1}^{C_{100}^{10}(C_9^1)^{90}(C_9^2+C_9^1)^{10}} \frac{1}{C_{100}^{10}(C_9^1)^{90}(C_9^2 + C_9^1)^{10}} log_2 \frac{1}{C_{100}^{10}(C_9^1)^{90}(C_9^2 + C_9^1)^{10}}$$

$$-\left(-\sum_{i=1}^{(C_9^1)^{100}} \frac{1}{(C_9^1)^{100}} log_2 \frac{1}{(C_9^1)^{100}}\right) = 67\ bits,$$

$$\Delta H_{conditionB} = -\sum_{i=1}^{C_{100}^{10}(C_9^1)^{100}} \frac{1}{C_{100}^{10}(C_9^1)^{100}} log_2 \frac{1}{C_{100}^{10}(C_9^1)^{100}} - \left(-\sum_{i=1}^{(C_9^1)^{100}} \frac{1}{(C_9^1)^{100}} log_2 \frac{1}{(C_9^1)^{100}}\right)$$

$$= 44\, bits\, .$$

Clearly, in a plastic network, the possibility of a bouton connecting to multiple dendrites dramatically increases the information entropy added by synaptic plasticity. The results hold when we scale up the model tenfold each time up to $10^6$ synapses (Figure 6H). The increase of information entropy values of 2 conditions under 5 model scales is shown in Table 4. Notably, whereas multi-dendritic MSB only slightly adds to the ISC in a static network (Figure 6G), the increase in the ISC for condition A by adding synapses is more than 50% (Figure 6H) higher than that of condition B in the plastic network. The relative advantage of multidendritic connectivity in $\Delta H$ scaled linearly with the network size (Figure 6I).

**Table 4. the increase of the ISC of a dynamic neural network**

| Model scale | $10^2$ | $10^3$ | $10^4$ | $10^5$ | $10^6$ |
|---|---|---|---|---|---|
| Condition A | 67 bits | 697 bits | 7,006 bits | 70,111 bits | 701,179 bits |
| Condition B | 44 bits | 464 bits | 4,684 bits | 46,892 bits | 468,986 bits |
